# The influence of polarized membrane ion carriers and extracellular electrical/pH gradients on cell ionic homeostasis and locomotion

**DOI:** 10.1101/2023.07.26.550658

**Authors:** Yizeng Li, Sean X. Sun

## Abstract

Anisotropic environmental signals or polarized membrane ion/solute carriers can generate spatially-varying intracellular gradients, leading to polarized cell dynamics. For example, directional migration of neutrophils, galvanotaxis of glioblastoma, and water flux in kidney cells, all result from the polarized distribution of membrane ion carriers and other intracellular components. The underlying physical mechanisms behind how polarized ion carriers interact with environmental signals are not well studied. Here, we use a physiologically-relevant, physics-based mathematical model to reveal how ion carriers generate intracellular ionic and voltage gradients. The model is able to discern the contribution of individual ion carriers to the intracellular pH gradient, electric potential, and water flux. We discover that an extracellular pH gradient leads to an intracellular pH gradient via chloride-bicarbonate exchangers, whereas an extracellular electric field leads to an intracellular electric potential gradient via passive potassium channels. In addition, the mechanical-biochemical coupling can modulate actin distribution and flow, and create biphasic dependence of the cell speed on water flux. Moreover, we find that F-actin interaction with NHE alone can generate cell movement, even when other ion carriers are not polarized. Taken together, the model shows the importance of cell ion dynamics in modulating cell migration and cytoskeletal dynamics.

## 1 Introduction

In cells, the spatial distributions of proteins and organelles are often not uniform. A polarized cell is essential in many (patho)physiological processes such as morphogenesis, immune response, nutrient delivery and filtration, signal transduction, and cancer metastasis [1,2]. Many studies have examined how extracellular biochemical signals generate cell polarization and establish intracellular gradients [3]. Cell polarity influences cell cytoskeletal structure, force distribution, morphology, and migration [4–9]. Recent works have shown that even in the absence of extracellular signals, the extracellular fluid influence cell polarity, giving a polarized distribution of ion channels, exchangers, and pumps. [10–13]. (Here, we will refer to these ion channels, exchangers, and pumps as membrane ion carriers.) Epithelial cells and cells in confinement display polarized distributions of ion carriers at the two ends of the cells without extracellular signal gradients [11,14–16]. In contrast, unconfined cells on two-dimensional substrates show a lesser degree of ion carrier polarization [17]. Polarized ion carriers can generate water fluxes across the cell membrane, contributing to processes such as cell shape change, epithelial fluid transport, and water-driven cell migration.

Since fluxes through ion carriers (and thus water) are affected by electric potential, any external electric fields will also generate intracellular ionic gradients. Moreover, fluxes of major ions such as sodium, potassium, and chloride are influenced by proton flux. Since the tumor microenvironment often has an elevated pH [18], and leukocytes [19, 20] and cancer cells [14–17] both utilize sodium-hydrogen exchangers to migrate, it is important to understand how environmental pH changes influence cell ionic homeostasis and migration. In this paper, we explore how extracellular electric potential and pH gradients affect intracellular ionic distribution through ion carriers using a theoretical model (Fig. 1A). We also examine how a polarized distribution of ion carriers can generate intracellular gradients without external signals (Fig. 1A). Since ions often interact strongly with proteins and genetic material, intracellular ionic gradients will contribute to intracellular protein biochemical gradients.

**Figure 1:**
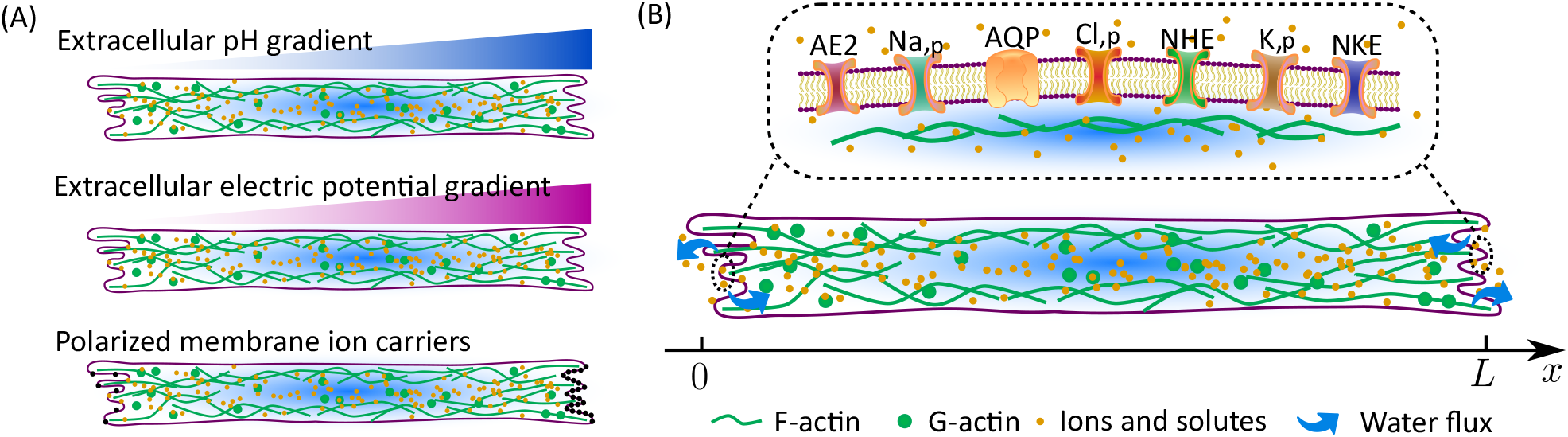
(A) The scenarios studied in this work. Cells are exposed to extracellular pH or electric potential gradients, and/or exhibiting polarized membrane ion carriers. (B) The elements included in the model. Na,_p_: passive sodium channel. K,_p_: passive potassium channel. Cl,_p_: passive chloride channel. NKE: sodium-potassium pump. NHE: sodium-hydrogen exchanger. AE2: chloride-bicarbonate exchanger. AQP: aquaporin. (Not to scale)

In addition to confinement, polarized ion carriers can occur through spatially varying F-actin assembly and disassembly, which not only regulate the distribution of ion carriers through vesicular trafficking but also through direct physical interaction with membrane-embedded proteins [21]. Our earlier work demonstrated that actin co-localizes with sodium-hydrogen exchangers in confined breast cancer cells [16] and with sodium-potassium pumps in kidney cells [11]. Polarization of ion carriers will, in turn, affect intracellular cell homeostasis and biochemical gradients, forming feedback loops.

A physiology-relevant, physics-based mathematical model has the potential to reveal cell migration mechanisms and provide unique insights into critical biophysical processes that are difficult to obtain from experiments. In this work, we use such a mathematical model to study the effects of (1) polarized ion carriers on cell migration and homeostasis, (2) polarized extracellular chemical-physical environment on cell polarization, and (3) actin-chemical coupling in cell polarization and water flux. The model predicts cell migration speed and cytoplasmic distributions of ionic concentrations, pH, and electric potential. We will also answer an interesting question: can there be just one single ion carrier responsible for sensing the cell microenvironment? The model predictions will have medical or therapeutic implications where cell dynamics are regulated by pH or electric potential [22, 23].

## 2 Method

The current theoretical model builds on our early work on steady-state cell migration in one-dimensional geometric confinement, but adds explicit analysis of pH and individual ionic fluxes. The model, when generalized to higher dimensions, can describe cell migration in any condition. In our results, we use *x* ∈ [0, *L*] to represent the cytoplasmic domain in the moving frame of the cell (Fig. 1B). The front of the cell is at *x* = *L* and the back is at *x* = 0. The model includes cytosol, F-actin, osmolarity, water flux [24], G-actin [25], charged solutes, pH, and electric potential [15,26]. The cytosol and extracellular water are connected through membrane water flux, *J*_water_. Depending on the cell morphology and extracellular physical environments, cells can migrate via either actin-driven or water-driven or both [14, 15, 17, 24, 27]. The amount of water contribution to the cell velocity was shown to depend on the coefficient of extracellular hydraulic resistance [24, 27]. In this work, we use a moderate hydraulic resistance so that a fraction (approximately 2.5%) of water flux converts to cell velocity. This choice is arbitrary because the hydraulic resistance is a free parameter. In real situations, the extracellular hydraulic resistance depends on geometry and fluid permeability of the environment, and can be higher or lower than the value we are using, corresponding to a larger or smaller contribution to water-driven cell velocity [24, 27].

In the cell, G-actin (concentration *θ*_*c*_) continuously polymerizes into F-actin (concentration *θ*_*n*_) through boundary actin polyermization fluxes at the cell front, *J*_actin_, and F-actin depolymerizes into G-actin in the cytoplasm with a rate, *γ* [25, 28]. The charged solutes considered in the model include sodium (Na^+^), potassium (K^+^), chloride (Cl^−^), hydrogen ion (H^+^), bicarbonate 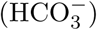, deprotonated buffer (Buf^−^), and non-permeable protein molecules (A^−^). To model ion fluxes across ion carries on the cell membrane, we consider passive sodium (Na_,p_), potassium (K_,p_), and chloride (Cl_,p_) channels; sodium-potassium pumps (NKE), sodium-hydrogen exchangers (NHE), and chloride-bicarbonate exchangers (AE2) (Fig. 1B).

We use *α*_*i*_ to represent the transport coefficient of each ion carrier, where *i*, such as *i* = NHE, is an index representing ion carriers. The transport coefficients depend on the activity and expression levels of the ion carriers. We quantify the degree of ion carrier polarization by the ratio 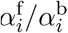, where the superscripts ‘f’ and ‘b’ refer to the quantities associated with the front and back of the one-dimensional cell, respectively. We take carrier polarization as given, meaning that we do not investigate how ion carriers are polarized via vesicular trafficking mediated by the cytoskeleton [29]. When varying the ratio 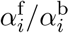, we keep the average transport coefficient fixed, i.e., 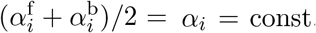. for each carrier. In this way, we focus on the effects of polarization, not the overall permeability of each ion carrier. Below we introduce the new model development beyond our previous work.

Many intracellular mechanical and biochemical processes are coupled to each other. Here we couple pH and NHE activity with the actin network using two known mechanisms. First, the rate of actin depolymerization, *γ*, depends on multiple mechanical and biochemical factors [30], including pH [31, 32]. Experimental data shows that higher pH leads to a higher rate of Cofilin-mediated actin depolymerization at the pointed end [31, 32]. Therefore, we let

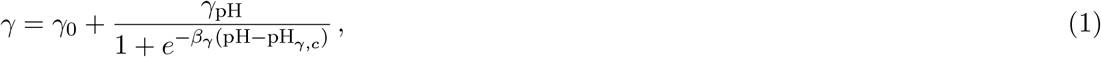

where pH is the local intracellular pH as a function of space. The intracellular pH is one of the unknowns to be solved by the model. *γ*_0_ is a baseline rate of actin depolymerization, and *γ*_pH_ is the coefficient of pH-dependent rate of actin depolymerization. *β*_*γ*_ and pH_*γ,c*_ are two constants characterizing how *γ* depends on pH.

Second, recent work showed that the activity of NHE correlates with F-actin density [16]. F-actin interacts with NHE through an actin binding protein, ezrin [21,33]. In this work, we will study, from a theoretical point of view, how this coupling affects cell ion homeostasis and migration. Since the membrane activity of NHE increases with increasing F-actin presence, we write the permeability coefficient of NHE as a function of F-actin concentration

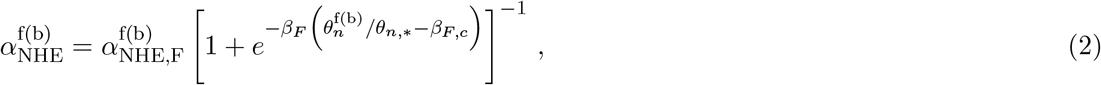

where 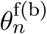 is the F-actin concentration evaluated at the front and back of the cell, respectively. 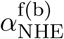 is the baseline permeability coefficient, which depends on the cell type and the expression level. *β*_*F*_, *β*_*F,c*_, and *θ*_*n,\**_ are constants. If we remove the actin-dependence of NHE activity, we let 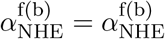, which can be different at the two ends of the cell.

A complete model description and parameters is provided in the Supplementary Materials. The chosen parameters represent typical mammalian tissue cells in physiological conditions, meaning these parameters generate pH, membrane potential, and ion concentrations within the expected physiological ranges. Cells in pathological conditions require a different parameter set. Given the potential variation from cell type to cell type, we do not focus on the particular results or numbers predicted by the model. Rather, we emphasize the mechanisms behind why the model predicts certain trends of cellular dynamics as functions of solute carrier polarizations, which can be observed in experiments via antibodies targeting specific carriers.

## 3 Results and Discussions

In our prior work, we showed how actin-driven and water-driven cell migration are modulated by the extracellular fluid environment, matrix adhesion, solute flux, and actin turnover [24, 25, 27, 28]. In particular, our models predict that polarized distributions of F-actin and actin retrograde flow towards the cell trailing edge as a result of the actin polymerization at the front. While F-actin polymerization is always polarized, the intracellular electro-neutral solute concentration, on the other hand, is evenly distributed if the solute flux is uniformly distributed across the cell membrane. This is still the case for the current model with multiple charged ions and ion carriers. We find that for a cell that resides in a uniform isotropic extracellular environment, without coupling between mechanical and biochemical processes, and without polarized ion carriers, every intracellular variable is uniform across the cell except for actin (Fig. S1A). Interesting spatial variations in ions and electrical potential occur only when the cell is polarized or there is coupling between mechanical and biochemical processes in the cytoplasm.

### 3.1 Polarization of ion carriers leads to intracellular gradients

Elongated cells have polarized distributions of ion carriers which enhance cell motility [15, 16]. This polarization also induces intracellular pH, electric potential, and osmotic gradients, which we can predict with our model. To begin with, we will remove the coupling between mechanics and chemistry; this coupling will be added later to examine the altered dynamics.

Frontal polarization of NHE plays critical roles in confined migration of human breast cancer and murine sarcoma cancer cells [14–16]. We use NHE as the first example to illustrate how ion carrier polarization leads to intracellular gradients. We vary the NHE polarization ratio from 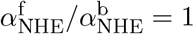, meaning non-polarized, to 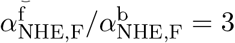, meaning a three-folds polarization to the cell front. In human breast cancer cells, 2.3 folds of front NHE polarization was observed [15].

Our model predicts that as the frontal polarization ratio of NHE increases, the intracellular ionic gradients increase (Fig. 2A). Higher pH is developed at the cell front than the cell back because NHE continuously removes hydrogen ions at the front. High pH leads to high bicarbonate concentration due to the bicarbonate-carbonic acid pair reaction. As a result, the spatial distributions of pH and bicarbonate concentration mirror each other. In addition, the frontal polarization of NHE brings in sodium at the cell front. Since NHE is electro-neutral, its polarization does not generate a significant intracellular electric potential gradient. Nor does it generate an intracellular cytosol pressure gradient or significantly modify the F-actin velocity and concentration (Fig. S1A).

**Figure 2:**
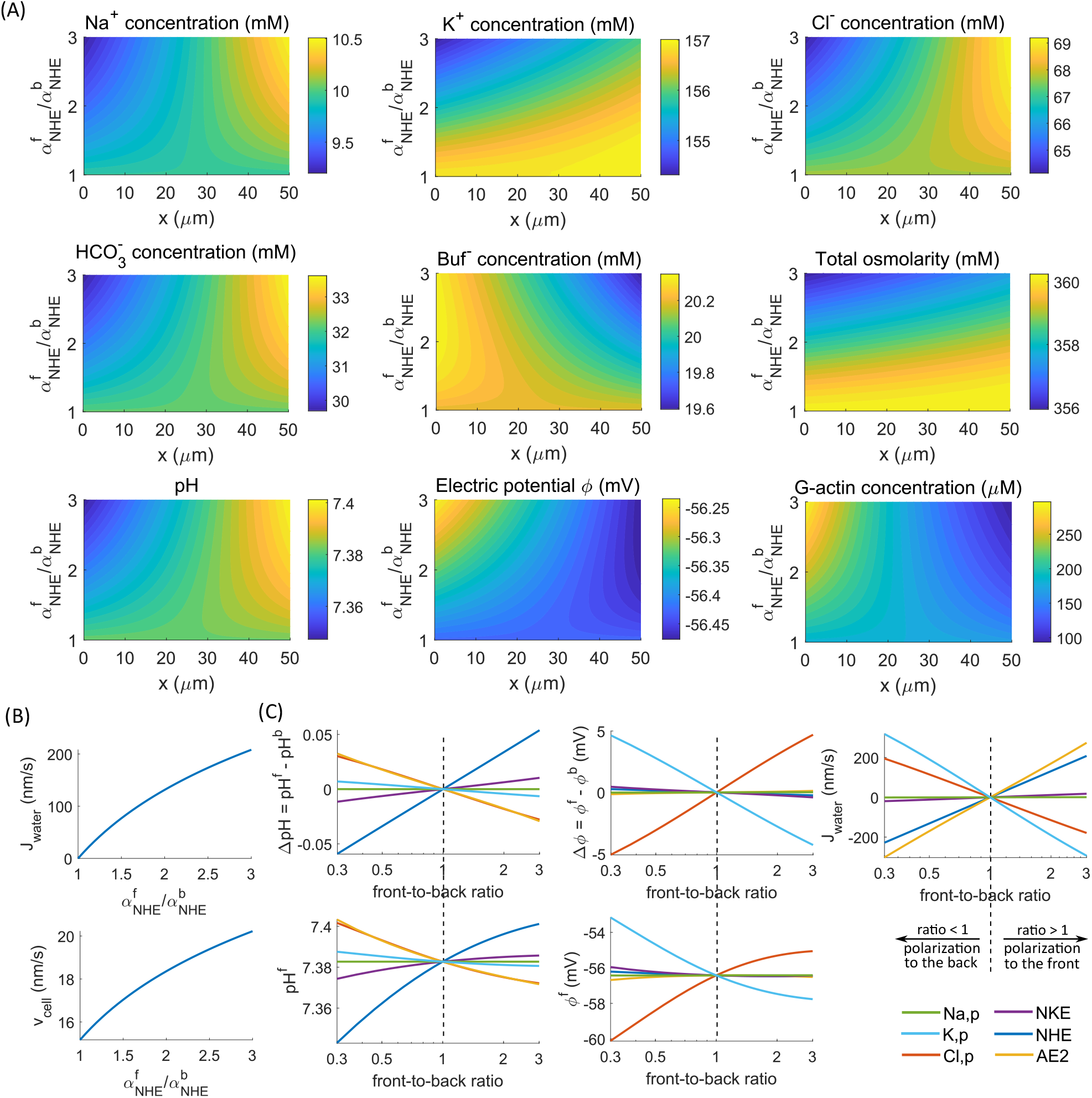
Model predictions on the effect of polarized ion carriers without coupling between mechanics and biochemistry. (A) Spatial distribution along the cell length (*x*) of intracellular variables for different degrees of frontal polarization of NHE 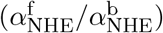. (B) Water flux (*J*_water_) and cell velocity (*v*_cell_) as functions of NHE polarization 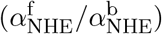. (C) Intracellular pH difference (ΔpH^f^ between the front and back intracellular pH), pH at the cell front (pH^f^), electric potential difference (Δ*ϕ* between the front and back intracellular electric potential), electric potential at the cell front (*ϕ*^f^), and water flux as functions of the polarization ratio of each ion carrier. A ratio equal to 1 means the ion carriers are not polarized. Each ion carrier is varied independently, while the other carriers remain non-polarized.

The combined gradients of all ion species generate an elevated osmolarity at the cell front (Fig. 2A), which drives water influx at the cell front (Fig. 2B). With contributions from water-driven cell migration, the cell velocity increases with NHE polarization as the water flux increases. This NHE-driven water influx is consistent with our understanding that high NHE expression level leads to cell swelling and water flow [21, 26, 34]. This water flux is predicted to drive G-actin towards the back of the cell, leading to a higher G-actin concentration at the cell back than the front (Fig. 2A). In summary, on the side where the NHE expression level is high, the overall osmolarity is also high but is less acidic. Water flows from the high NHE side to the low side.

Not all ion carriers are equally effective in generating intracellular ion gradients. Our model predicts that the polarization of NHE, AE2, or the passive chloride channel can modulate the pH gradient, but not other ion carriers (Fig. 2C, ΔpH). NHE and AE2 involve hydrogen and bicarbonate ions, respectively, and thus directly impact pH. The passive chloride channel affects chloride concentration, which interacts with AE2, thus indirectly affecting pH. Frontal polarization of NHE increases the pH value at the front, whereas AE2 and the passive potassium channel decrease it (Fig. 2C, pH^f^). However, due to their electro-neutrality, NHE and AE2 are inefficient in modulating the intracellular electric potential (Fig. 2C, *ϕ*^f^ and Δ*ϕ*). In contrast, the polarization of the passive potassium and chloride channels can affect the electric field significantly because of their high flux and electrogenic nature.

Our model predicts that ion carriers that are effective in generating pH or electric potential gradients are also effective in generating water flux (Fig. 2C, *J*_water_). Frontal polarization of NHE and AE2 leads to water influx at the front (and thus efflux at the back). In contrast, the frontal polarization of the passive potassium and chloride channels leads to water efflux at the front (and thus influx at the back). This model prediction is consistent with experimental observations: frontal polarization of NHE1 and back polarization of SWELL1, a type of passive chloride channel, collectively drive cell migration [15]. AE2 and the passive chloride channel generate intracellular pH gradients along their direction of polarization but create water flux in the opposite direction.

The polarization of NKE, one of the essential ion carriers in maintaining physiological functions, seems to have little effect on intracellular pH and electric potential compared to other ion carriers (Fig. 2C). This does not mean that NKE does not matter. The model predicts that NKE polarization ratio needs to be significantly larger to see differences in the pH gradient, electric potential, and water flux (Fig. S1B). When NKE is highly polarized to the back, water flows out of the cell at the back. This directional water flux has also been demonstrated in experiments where kidney cells pump water from the apical to the basal side [11].

### 3.2 Extracellular pH polarization leads to intracellular pH gradient via AE2

Extracellular pH gradients can occur *in vivo* in multiple scenarios, such as from solute gradients generated by neighboring cells, fluid circulation in organs, the tumor microenvironment, or during immune response [35]. At the same time, cells migrating in this environment may have polarized ion carriers. We investigate how extracellular pH gradients interact with polarized ion carriers during migration. In the model, we increase the extracellular pH at the front of the cell, 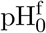, while keeping the pH at the back, 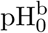, fixed at 7.42. The increase of 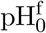 is obtained by increasing the extracellular bicarbonate concentration at the front.

In cells without ion carrier polarization, the intracellular pH gradient follows the extracellular one (Fig. 3A). This is because increased extracellular bicarbonate concentration at the front leads to less bicarbonate efflux through AE2, and thus increases intracellular bicarbonate concentration at the cell front (Fig. 3A). For this reason, we hypothesize that AE2 is the first responder to the extracellular pH change, and other ion carriers follow the AE2 change instead of following the extracellular cues. We will test this hypothesis later. Cells have abilities to cushion environmental stress, so the intracellular pH gradient is much smaller than the extracellular one, as predicted by the model. The pH gradient has little effect on the intracellular electric potential (Fig. S2).

**Figure 3:**
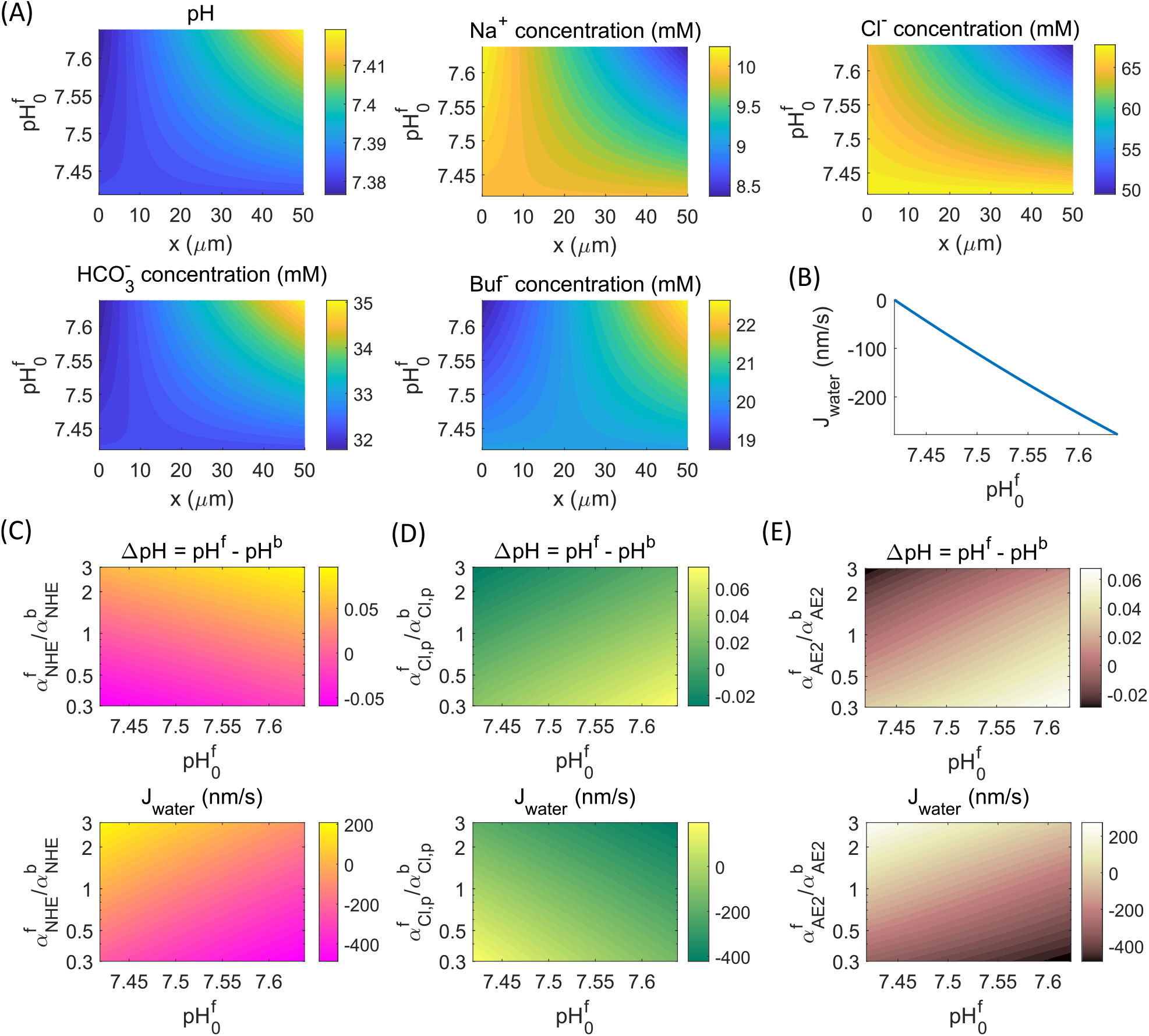
Model prediction where there is an extracellular pH increase (more alkaline) at the cell front while the pH at the back is fixed. Extracellular pH change is obtained by changing the concentration of extracellular bicarbonate. Before polarization, 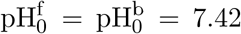. After polarization, 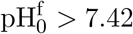 and the value varies. No coupling between mechanics and chemistry. (A) Spatial distribution (*x*) of intracellular pH and some ion concentrations as functions of extracellular pH at the cell front. (B) Water flux at the cell front as a function of extracellular pH at the cell front. (C)-(E) The intracellular pH difference (defined as the pH at the cell front minus the back) and water flux at the cell front as functions of ion carrier polarization (NHE in panel C, passive chloride channel in panel D, and AE2 in panel E) and extracellular pH at the cell front.

The spatial gradient of the bicarbonate concentration mirrors that of pH (Fig. 3A), as we have seen in the last section (Fig. 2A). Although a frontal polarization of NHE and an extracellular pH increase at the front generate same intracellular pH and bicarbonate gradients, the other ionic concentration fields are different between these two scenarios (Figs. 3A and 2A). In addition, with increased extracellular pH at the cell front, the model predicts water efflux at the front (Fig. 3B), as opposed to influx with NHE polarization (Fig. 2B). These phenomena suggest that the intracellular pH gradient alone does not determine the distribution of ions and the direction of water flux.

We thus expect that the combination of extracellular pH and NHE polarizations generate interesting patterns. Indeed, the model predicts that increasing extracellular pH at the cell front while polarizing NHE towards the back cancels the intracellular pH gradient, but polarizing NHE towards the cell front enhances the positive intracellular pH gradient brought by the high front extracellular pH (Fig. 3C). The water flux follows an opposite pattern such that a frontal polarization of NHE cancels the water flux generated by front extracellular pH increase.

Likewise, the combination of an extracellular pH gradient with the polarization of the passive chloride channel (Fig. 3D) or AE2 (Fig. 3E) has different effects on generating intracellular pH gradient and water flux. Cells in general can have multiple ion carriers polarized at the same time. Since ion fluxes are additive, the effects from these polarizations are also additive. For example, a frontal polarization of NHE and a back polarization of the passive chloride channels mutually increase water influx at the cell front, whereas frontal polarization of both NHE and the passive chloride channels results in no water flux or even water efflux at the cell front [15].

To test our hypothesis that AE2 is *the* carrier that responds to the extracellular pH change, we vary the AE2 permeability at the cell front while keeping it unchanged at the back and observe the cell responses in the absence of extracellular pH polarization. The model indeed predicts that a decrease of AE2 permeability at the cell front produces the same intracellular gradients of all field variables as the case of increasing the extracellular pH at the cell front (compare Fig. S3 with Fig. S2). No other ion carriers are able to achieve this. Therefore, from the cell homeostasis regulation perspective, a higher extracellular pH at the cell front (without regulating AE2 activities) is equivalent to reducing AE2 activity at the cell front (without polarizing the extracellular pH).

### 3.3 Extracellular electrical potential gradients leads to intracellular gradients via passive potassium channels

We next study the effect of extracellular electric potential gradients by increasing the front potential, 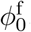, from 0 to 30 mV while keeping 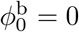 at the back. In cells without ion carrier polarization, the intracellular electric potential gradient is predicted to follow the extracellular one (Fig. 4A) but its gradient is smaller. The electric potential gradient has little effect on pH (Fig. S4).

**Figure 4:**
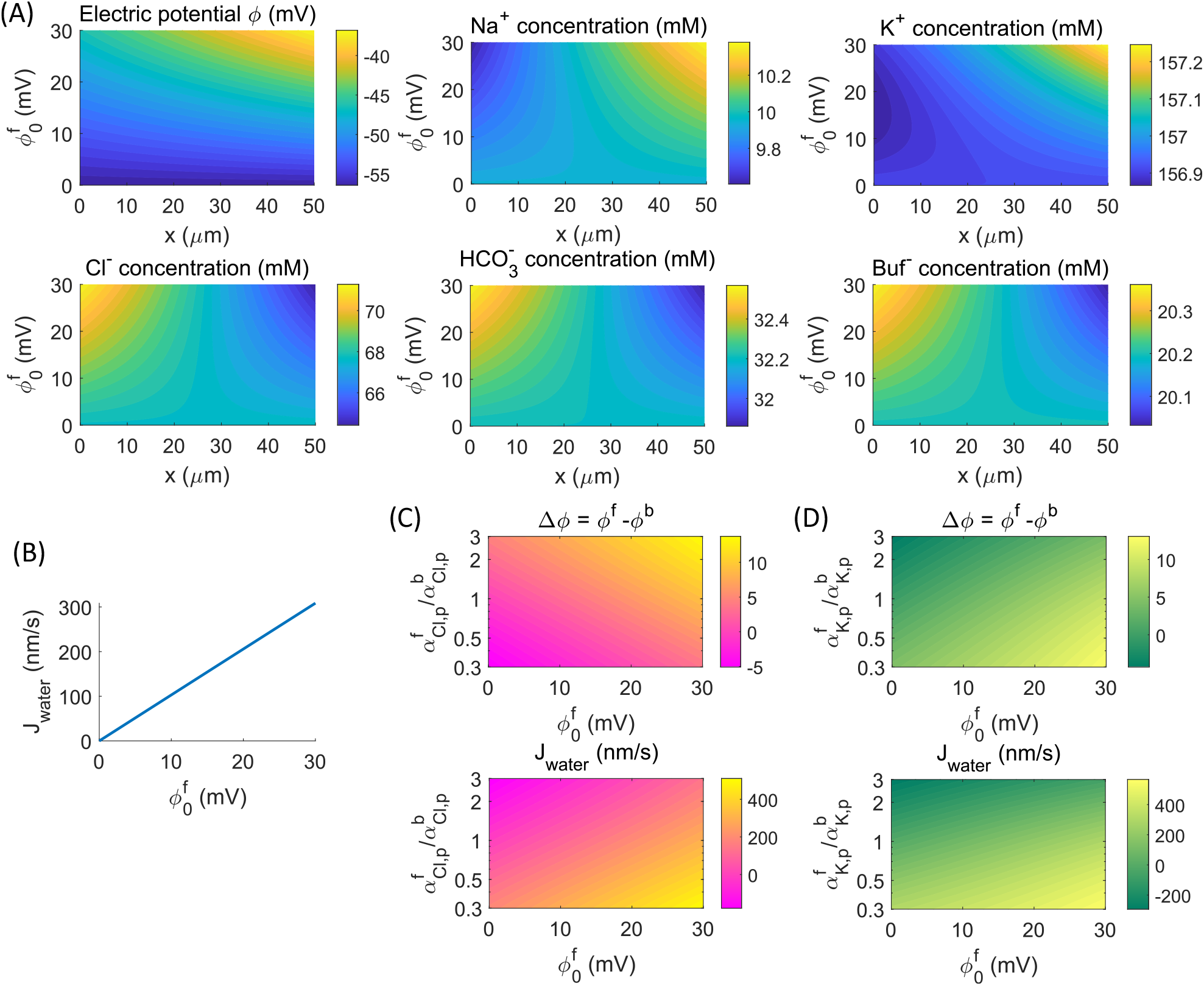
Model predictions on the effect of extracellular electric potential increase at the cell front while the potential at the back is fixed. Before polarization, 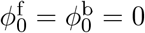. After polarization, 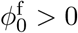 and the value varies. No coupling between mechanics and chemistry. (A) Spatial distribution (*x*) of intracellular electric potential and some ion concentrations as functions of extracellular electric potential at the cell front. (B) Water flux at the cell front as a function of extracellular electric potential at the cell front. (C)-(D) The intracellular electric potential difference (defined as the potential at the cell front minus the back) and water flux at the cell front as functions of ion carrier polarization (passive chloride channel in panel C and passive potassium channel in panel D) and extracellular electric potential at the cell front.

A positive extracellular electric potential at the front leads to reduced potassium efflux from the passive potassium channel, generating a relatively high intracellular potassium concentration at the cell front (Fig. 4A). Fluxes through other ion carriers adjust as well, creating intracellular gradients for all ions. However, the gradients of ion concentrations from external electric potential gradients are smaller than those from the pH gradient (compare Figs. 4A and 3A), although the magnitude of water fluxes from the two cases are comparable (Figs. 4B and 3B).

The passive chloride and potassium channels are the only two ion carriers whose polarization is able to generate intracellular non-trivial electric potential gradient (Fig. 2C). When these two channels are polarized in the presence of an extracellular electric potential gradient, various patterns of potential gradients and water flux occur (Fig. 4C,D). For example, a frontal polarization of the chloride channel increases the intracellular electric potential generated by the extracellular potential but reduces or even reverses the water flux. Our model suggests that cells *in vivo* may develop ion carrier polarization to enhance or counter-act environmental electrical signals.

Since passive chloride channels generate an intracellular pH gradient (Fig. 2C), but there is almost no intracellular pH gradient under extracellular electric potential polarization (Fig. S4), we thus hypothesize that the passive potassium channel is the agent that responds to the extracellular electric potential signal. We test this hypothesis by varying the permeability of the passive potassium channel at the cell front in the absence of extracellular electric potential polarization. The model predicts that reducing front potassium channel activities is equivalent to increasing the extracellular front electric potential (compare Fig. S5 with Fig. S4). All spatial fields have the same directions of gradients, and the water fluxes follow the same direction under these two cases. No other ion carriers can achieve this correspondence.

### 3.4 pH-triggered actin depolymerization can modulate F-actin distribution

In the above discussion, we used a decoupled model between actin and pH (*γ* = *γ*_0_ = const. and *γ*_pH_ = 0 in Eq. 1) to analyze how each ion carrier impacts cell homeostasis. In this section, we will study the effect of pH-dependent actin depolymerization on the F-actin distribution. Recall that polarization of extracellular pH leads to membrane water flux (Fig. 3B). Water flux generates cytosol flow, which impacts actin distribution in two ways.

The first effect comes from the convection of G-actin by the cytosol flow. When the cytosol flows from the back to the front of the cell, which happens when 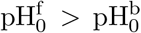, it carries G-actin and polarizes it towards the front (Fig. 5A). When the opposite occurs, G-actin polarizes to the back (Fig. 5B). The second effect is on F-actin concentration influenced by G-actin. In the simplest case with no environment or ion carrier polarization, F-actin always concentrates at the cell front where actin polymerization occurs (Fig. S1A). This frontal F-actin polarization can be critical in actin-driven cell migration [25, 28, 36]. However, when G-actin polarizes towards the back, which occurs when cytosol flows from the front to the back, F-actin can redistribute (Fig. 5B).

**Figure 5:**
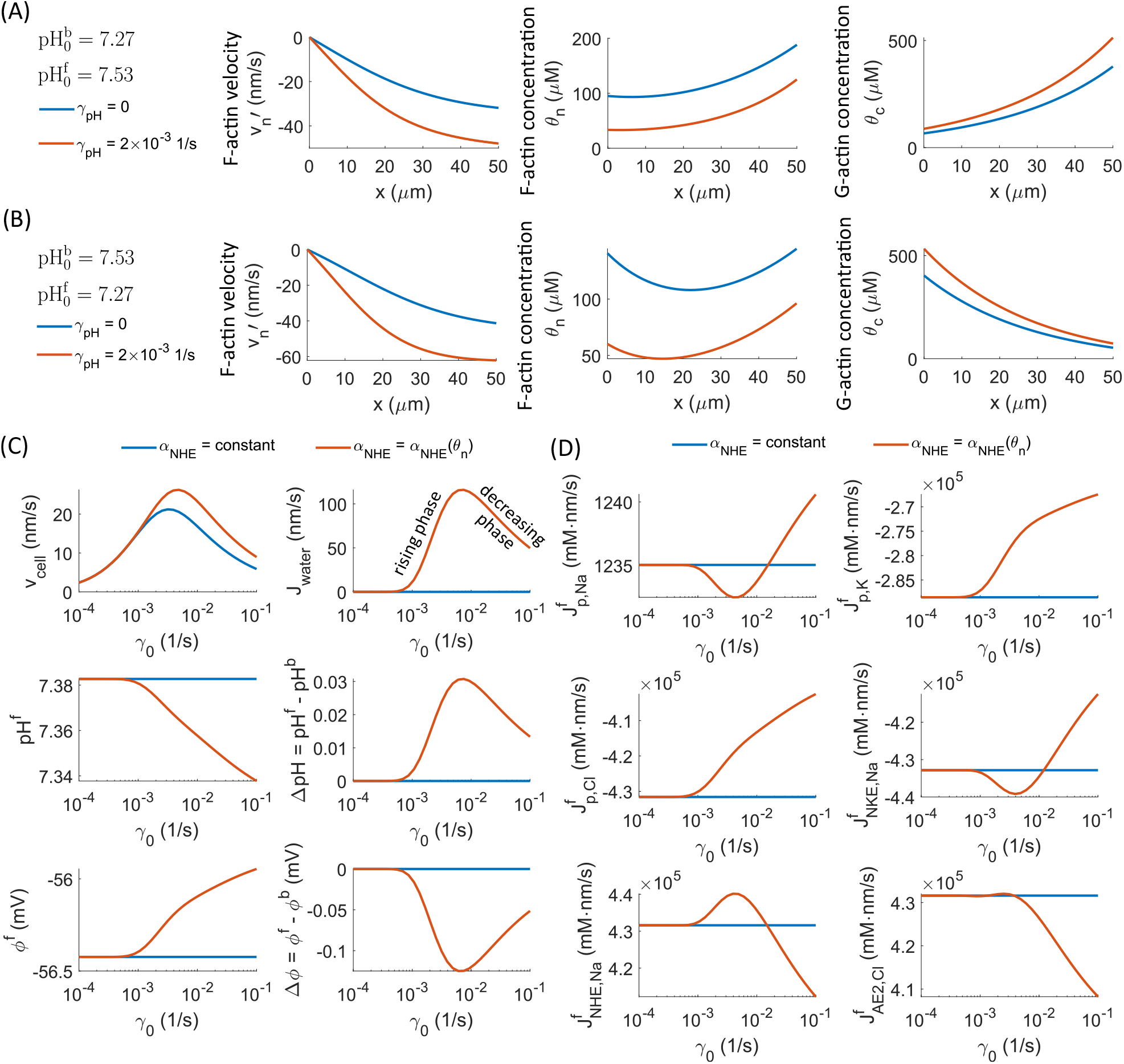
Effects of coupling among actin depolymerization, pH, and NHE activities. (A) Actin distribution in a polarized extracellular pH environment. 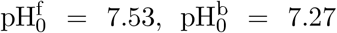. 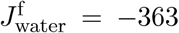 nm/s from model calculation. (B) Actin distribution in a polarized extracellular pH environment. 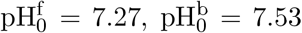. 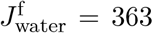 3 nm/s from model calculation. (C) Cell velocity and homeostasis in a non-polarized environment as functions of a constant rate of actin depolymerization for different NHE permeability choices. (D) Fluxes through ion carriers in a non-polarized environment as functions of a constant rate of actin depolymerization for different choices of NHE permeability.

The F-actin distribution can revert when the rate of actin depolymerization, *γ*, increases with pH by cofilin [31, 32]. Since the intracellular pH gradient follows the extracellular one, so does the actin depolymerization rate. When 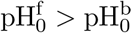, a higher rate of actin depolymerization at the cell front does not change the polarity of F-actin (Fig. 5A). When 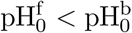, the higher rate of actin depolymerization at the cell back disassembles F-actin, leading to a frontal polarization of the cell (Fig. 5B) similar to that in the control case (Fig. S1).

The model suggests that pH can modulate F-actin distribution through pH-induced cytosol flow, pH-dependent actin depolymerizatoin, or both. Since actin distribution is critical to cell motility, modulating intracellular dynamics, and force distribution, the coupling between pH and actin can be one of the mechanisms to ensure proper cell function when there is environmental pH shocks.

### 3.5 Actin-generated NHE polarization generates biphasic water flux and intracellular pH gradient

We have so far prescribed ion carriers’ polarization by setting their front-to-back polarization ratios as parameters. In this section, we consider the case where NHE polarization is determined by F-actin [16, 21], i.e., *α*_NHE_ = *α*_NHE_(*θ*_*n*_) (Eq. 2), instead of being prescribed. In order to focus on the effect of actin-induced NHE polarization, we let *γ* = *γ*_0_ = const., and the other ion carriers and the extracellular environment are non-polarized.

Without actin-induced NHE polarization, there is no water flux nor intracellular ionic gradients regardless of the rate of actin depolymerization, *γ*, in a non-polarized cell (Fig. 5C and D). The cell velocity is a biphasic function of *γ* because of an actin-flow to actin-distribution transition [28]. The rate of actin depolymerization, *γ*, reduces the average F-actin concentration across the cell and establishes a high F-actin polarization ratio, i.e., 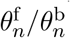 increases with *γ* [28]. After implementing actin-induced NHE polarization where the permeabilities 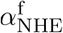 and 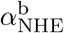 follow 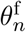 and 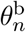 NHE polarizes towards the cell front as the rate of actin depolymerization, *γ*, increases. The model predicts a biphasic water flux as a function of actin depolymerization rate, *γ* (Fig. 5C). The rising phase in the water flux comes from the increasing ratio of NHE polarization as *γ* increases. The decreasing phase is due to reduced NHE flux as F-actin concentration drops. In this case, even if the NHE polarization ratio is very large, the low NHE flux cannot establish a large osmotic gradient. Biphasic water flux contributes to biphasic cell velocity (Fig. 5C).

The three ion carriers that involve sodium ions, NHE, NKE, and the passive sodium channel, also show biphasic fluxes as *γ* increases. Ion carriers that do not involve sodium are monotonic in *γ*, except for AE2, where a small bump appears due to its linkage to pH (Fig. 5D). With the actin-NHE coupling, our model indicates that F-actin alone can generate water flux and ion dynamics without prescribing polarization for other membrane ion carriers. This result is surprising and suggest that the actin-NHE interaction is the central element in establishing cell movement. The gradients of intracellular pH and electric potential also show biphasic relations with the rate of actin depolymerization (Fig. 5C, ΔpH and Δ*ϕ*) but not the baseline pH and electric potential values (Fig. 5C, pH^f^ and *ϕ*^f^). This difference suggests that pH and electric potential gradients are influenced by both the NHE polarization ratio and absolute permeability. In contrast, pH and electric potential values are only affected by NHE absolute flux. The decreasing pH with increasing *γ* indicates that cells will not further elevate actin depolymerization through pH-*γ* coupling.

## 4 Conclusions

In this work, we used a theoretical model to study the impact of ion carriers and extracellular environment gradients on cell ion homeostasis, migration, and actin network distribution. Each ion carrier plays a specific role in maintaining cell homeostasis by communicating information between the inside and the outside. Some ion carriers modulate pH, some modulate electric potential, and some do both. The model predicts that pH and electric potential are uncoupled, suggesting that cells can maintain homeostasis of one field when the other field is perturbed. Our results show that, within the ion carriers included in this work, extracellular pH gradients lead to intracellular pH gradient via chloride-bicarbonate exchangers, whereas extracellular electric potential polarization leads to intracellular electric potential gradient via passive potassium channels. The magnitudes of intracellular gradients are smaller than the extracellular ones, indicating that cells can cushion environmental changes through negative feedback.

Cells develop intracellular pH, electric potential, and ion concentration gradients when membrane ion carriers are polarized. The magnitudes of these gradients depend on the ratio of carrier polarization. The types of membrane ion carriers also play a major role in determining the gradients. Given the same polarization ratio, NHE, AE2, and the passive chloride and potassium channels are more effective in creating intracellular biochemical gradients than NKE and the passive sodium channel. When ion carrier polarization is coupled with extracellular polarization, various combinations of intracellular electric potential gradient, pH gradient, and water flux can occur. It is possible to create an intracellular gradient without generating water flux, or generate water flux without creating an intracellular gradient. When extracellular hydraulic resistance is high, water flux will increase the cell migrations speed.

The coupling between the actin network and ion carriers introduces more complexity and increases the diversity of behaviors we observe. Our model predicts that the polarization of F-actin concentration can change when an intracellular pH gradient develops. The distribution of F-actin is related to many intracellular processes, such as force distribution, morphology, nucleus anchoring, nuclear envelope stress, and intracellular trafficking. In addition to the known impact of pH on cell biochemistry, the model suggests that pH also has the potential to indirectly affect cell dynamics through a redistribution of the actin network.

The structural linkage between F-actin and NHE through ezrin suggests that the polarization of F-actin establishes NHE polarization. The polarization of F-actin comes from actin polymerization, and the polarization of NHE leads to water flux. As a result, actin-driven cell migration and water-driven cell migration are intrinsically coupled together. In two-dimensional cell migration, where water-driven cell migration is not applicable due to the low extracellular hydraulic resistance, we can still expect intracellular cytosol flow generated by transmembrane water flux. Such cytosol flow has the potential to provide effective convection for intracellular signaling molecules. Moreover, our model predicts that F-actin alone can generate water flux through NHE, even when other membrane ion carriers are not polarized. This result is surprising and suggest that the actin-NHE interaction is the central element in establishing cell movement.

To summarize, our model predicts that membrane ion carriers and environmental polarizations can generate intracellular osmotic, electric potential, pH, and actin gradients, which have implications on intracellular dynamics and signaling. The coupling between biochemistry and mechanics further increases the cells’ adaptability to internal or external perturbations. The future development of understanding on cell biology requires integrated views of biochemical and physical processes.

## Acknowledgment

Yizeng Li is supported by NSF 2303648. Sean X Sun is supported by NIH R01GM134542. The opinions, findings, and conclusions, or recommendations expressed are those of the authors and do not necessarily reflect the views of any of the funding agencies.

