## Supplementary Materials for "The influence of polarized membrane ion carriers and extracellular electrical/pH gradients on cell ionic homeostasis and locomotion"

### 1 Model Details

#### 1.1 Model overview

Here we describe a steady-state cell migration model with the consideration of cytosol, F-actin, G-actin, charged ions, pH, and voltage. The model is applicable to confined cell migration, either in a narrow channel or three-dimensional extracellular matrix, where cells are elongated and migrate along one direction, which we will denote as the $x$-direction. All field variables are thus functions of $x$. We neglect the changes of variables along the directions perpendicular to the cell migration. In many cases, space exists between the nucleus and the cell lateral wall, so the nucleus does not significantly block the material transportation from one end of the cell to the other [1–4]. For this reason, we do not explicitly model the nucleus. The effect of the nucleus can be incorporated into the effective diffusion coefficients of the ions and proteins, if desirable. It is also possible that the nucleus separates a cell's front and back compartments and develops pressure differences within the cell [5, 6]. Our model does not include this scenario.

The cell front is defined by where actin polymerization happens, which is also the direction of cell migration without a strong water flow perturbation. We will use superscripts 'f" and 'b' throughout the paper to represent quantities associated with the front and back ends of the cell, respectively. Under steady state, the cell velocity, $v_{\text{cell}} = \dot{x}^{\text{f}}(t) = \dot{x}^{\text{b}}(t)$, and the cell length, $x^{\text{f}}(t) - x^{\text{b}}(t) = L$, are constants. Quantities associated with the extracellular environment will be denoted by a subscript '0'. In this work, we consider a more involved model compared to our early works [3, 4, 7–10]. This model emphasizes the coupling among the mechanical, electrical, and chemical properties of a cell and how the polarization of one aspect of the cell affects the other properties and cell migration.

The primary variables, or unknowns to be solved, of the model are

$$\mathbf{X} = \{p_c(x), v_c, v_n(x), \theta_n(x), \theta_c(x), c_{\text{Na}}(x), c_{\text{K}}(x), c_{\text{Cl}}(x), \text{pH}(x), c_A(x), c_{\text{Buf}}(x), \phi(x), v_{\text{cell}}\}^T \ , \quad \text{(S1)}$$

*
†

$p_c(x)$: hydro-static pressure of the cytosol; unit: Pa.
$v_c$: velocity of the cytosol; unit: nm/s.
$v_n(x)$: velocity of F-actin; unit: nm/s.
$\theta_n(x)$: concentration of actin in the filamentous form (F-actin); unit: mM.
$\theta_c(x)$: concentration of actin in the monomeric form (G-actin); unit: mM.
$c_{\text{Na}}(x)$: concentration of $\text{Na}^+$ in the cytosol; unit: mM.
$c_{\text{K}}(x)$: concentration of $\text{K}^+$ in the cytosol; unit: mM.
$c_{\text{Cl}}(x)$: concentration of $\text{Cl}^-$ in the cytosol; unit: mM.
pH$(x)$: intracellular pH.
$c_A(x)$: concentration of non-permeable proteins, $\text{A}^-$, in the cytosol; unit: mM.
$c_{\text{Buf}}(x)$: concentration of deprotonated buffer ions, $\text{Buf}^-$, in the cytosol; unit: mM.
$\phi(x)$: intracellular electric potential; unit: mV.
$v_{\text{cell}}$: steady-state cell migration velocity; unit: nm/s.
These variables will be solved through a set of coupled non-linear equations, which we will describe in detail below. All other variables can be derived from these primary variables. This model does not include myosin dynamics but can be expanded to include it if necessary, as we have developed in our prior work [11].

##### 1.2 Cytosol mechanics

The cytosol is a water-like fluid that exchanges with the extracellular water through membrane water flux. The relative motion between the cytosol and the actin network generates an interface stress proportional to the velocity difference of the two phases, i.e., $\eta\theta_n(v_c - v_n)$, where $\eta$ is the coefficient of interfacial friction between the actin-network phase and the cytosol phase. When the interface stress is significantly larger than the cytosol viscous shear stress, the viscous shear stress can be neglected from the model. In this case, the pressure gradient of the cytosol balances the interfacial stress. The force and mass balances of the cytosol are

$$-\frac{dp}{dx} - \eta\theta_n(v_c - v_n) = 0\,, \quad \frac{dv_c}{dx} = 0\,. \tag{S2}$$

The mass balance in the one-dimensional space, $dv_c/dx = 0$, suggests that $v_c(x) = v_c = \text{const}$. The flux boundary conditions for cytosol are

$$v_c - v_{\text{cell}} = -J^{\text{f}}_{\text{water}}\,, \quad \text{at } x = x^{\text{f}}\,; \quad v_c - v_{\text{cell}} = J^{\text{b}}_{\text{water}}\,, \quad \text{at } x = x^{\text{b}}, \tag{S3}$$

where $J_{\text{water}}$ is the water influx across the cell membrane. Membrane water flux is driven by the combined hydrostatic and osmotic pressure difference of water across the cell membrane [12],

$$J^{\text{f(b)}}_{\text{water}} = -\alpha^{\text{f(b)}}\left[\left(p^{\text{f(b)}} - p^{\text{f(b)}}_*\right) - RT\left(c^{\text{f(b)}} - c^{\text{f(b)}}_0\right)\right]\,, \tag{S4}$$

where $\alpha$ is the permeability coefficient of water, $c$ is the total concentration of all ion species (will be discussed later), $R$ is the gas constant, and $T$ is the absolute temperature. Due to the hydraulic resistance, the fluid pressure exerted on the outside of the cell, $p_*$, differs from the fluid pressure at infinity, $p_\infty$. $p_*$ can be solved as

$$p^{\text{f}}_* = p^{\text{f}}_\infty + d^{\text{f}}_g(v_{\text{cell}} - J^{\text{f}}_{\text{water}})\,, \quad p^{\text{b}}_* = p^{\text{b}}_\infty - d^{\text{b}}_g(v_{\text{cell}} + J^{\text{b}}_{\text{water}})\,, \tag{S5}$$

where $d_g$ is the coefficient of external hydraulic resistance, which depends on the extracellular geometry and fluid viscosity. Approximated analytical expressions of $d_g$ in the one-dimensional and three-dimensional spaces have been derived and computed in our prior work [7, 13]. In this work, we will treat $d_g$ as a parameter. This coefficient of external hydraulic resistance determines whether cells can use water flux to drive cell migration. A large $d_g$ enables cells to convert water flux into motility, whereas cells in a low $d_g$ environment can only migrate via actomyosin dynamics [7,8].

##### 1.3 Actin network mechanics

The F-actin forms a fluid-like actin network. The existence of the actin filament generates passive swelling stress, $\sigma_n$, which can be modeled by a linear constitutive relation, $\sigma_n = k_{\sigma_n}\theta_n$, where $k_{\sigma_n}$ is the coefficient of actin swelling. A more involved constitutive relation from polymer physics can also be used [8]. Although we do not explicitly model the myosin in this work, the myosin contractile stress, $\sigma_a$, can still be included as a parameter if necessary. In this case, the total stress in the actin network will be $\sigma = \sigma_n - \sigma_a$. A model including myosin contractility can be found in our prior work [11].

The actin network connects to the extracellular matrix via focal adhesions [14]. As cells migrate, focal adhesions exert an effective body force on the actin network in the opposite direction of the actin flow, $v_n$. The magnitude of the body force depends on the magnitude of the actin flow and the distribution of the actin network. It can be modeled by $\eta_{\text{st}}\theta_n v_n$ [8,9], where $\eta_{\text{st}}$ is the coefficient of focal adhesion, which depends on the substrate stiffness [15,16] and the size [17] and density [18] of adhesions. In this work, we treat $\eta_{\text{st}}$ as a parameter and do not include the dynamics of focal adhesion [19].

The interfacial friction exerted on the cytosol also applies to the actin network. Putting all the forces together, we can write the force balance of the actin network,

$$-\frac{d\sigma}{dx} - \eta\theta_n(v_n - v_c) - \eta_{\text{st}}\theta_n v_n = 0\,. \tag{S6}$$

Since the actin network stress appears in the spatial derivative, the constant parameter of the myosin contractile stress, $\sigma_a$, does not influence the force balance of the actin network. For this reason, the current paper does not emphasize the role of myosin.

##### 1.4 F-actin and G-actin exchange

Actin polymerization and depolymerization are essential processes in actin-driven cell migration. Polymerization typically occurs at the cell's front edge, whereas depolymerization occurs throughout the cytoplasm. Depolymerization can be considered a sink for F-actin but a source for G-actin. The amount of actin depolymerization per unit of time depends on the concentration of F-actin, which can be represented by $\gamma\theta_n$, where $\gamma$ is the rate of actin depolymerization. Thus the material balance for F-actin and G-actin are

$$\begin{aligned} &\frac{d}{dx}(\theta_n v_n) = -\gamma\theta_n\,, \\ &\frac{d}{dx}(\theta_c v_c) = D_{\theta_c}\frac{d^2}{dx^2}\theta_c + \gamma\theta_n\,, \end{aligned} \tag{S7}$$

where $D_{\theta_c}$ is the diffusion coefficient of G-actin in the cytosol.

The contribution of actin polymerization is modeled through flux boundary conditions of Eq. S7.

At the front of the cell, the flux boundary conditions for F-actin and G-actin are

$$\theta_n(v_n - v_{\text{cell}})\big|_{x=x^{\text{f}}} = -J^{\text{f}}_{\text{actin}}\,, \quad \left[-D_{\theta_c}\frac{d\theta_c}{dx} + \theta_c(v_c - v_{\text{cell}})\right]\big|_{x=x^{\text{f}}} = J^{\text{f}}_{\text{actin}}\,, \tag{S8}$$

where $J^{\text{f}}_{\text{actin}}$ is the rate of actin polymerization. We assume that the rate of actin polymerization increases with the concentration of G-actin, $\theta_c$, and saturates when $\theta_c$ is large. Therefore, $J^{\text{f}}_{\text{actin}}$ takes the form

$$J^{\text{f}}_{\text{actin}} = J_a\,\frac{\theta^{\text{f}}_c}{\theta_{c,c} + \theta^{\text{f}}_c}\,, \tag{S9}$$

where $J_a$ is the coefficient of actin polymerization and $\theta_{c,c}$ is a constant. $\theta^{\text{f}}_c$ is the concentration of G-actin at the front of the cell, i.e., $\theta^{\text{f}}_c = \theta_c\big|_{x=x^{\text{f}}}$. Since there is no polymerization at the back of the cell, the fluxes for F-actin and G-actin are zero at $x = x^{\text{b}}$.

Within the time scale of consideration, the total amount of actin is conserved so that the average concentration of actin, $\theta_*$, should be a constant, i.e.,

$$\int_{x^{\text{b}}}^{x^{\text{f}}} (\theta_n + \theta_c)dx = L\theta_*\,. \tag{S10}$$

In the model, $\theta_*$ is prescribed but $\theta_c$ and $\theta_n$ are solved.

#### 1.5 Solute species and osmolarity

The boundary condition of the cytosol involves membrane water flux (Eq. S4) as a function of the total solute concentration. The solutes are essential because they determine the osmotic pressure and specify the pH and electric potential. We include in the model the ion species most abundant in cells (such as sodium, potassium, and chloride) and are crucial to studying the pH (i.e., hydrogen) and electric potential. We cannot neglect the concentration of bicarbonate if hydrogen is included because of the bicarbonate-carbonic acid pair reaction.

Within the cell, we consider the following solute species: $Na^+$, $K^+$, $Cl^-$, $H^+$, $HCO_3^-$, $A^-$, $Buf^-$, and HBuf. $A^-$ is the non-permeable large molecules or proteins, which we assume has valence $-1$, $Buf^-$ is a generic deprotonated buffer species, and HBuf is the corresponding protonated buffer species. So the intracellular solute concentration space is $c_i = \{c_{\text{Na}}, c_{\text{K}}, c_{\text{Cl}}, c_{\text{H}}, c_{\text{HCO}_3}, c_{\text{A}}, c_{\text{Buf}}, c_{\text{HBuf}}\}^T$. Outside of the cell, we neglect the buffer solution but include an electro-neutral, non-permeable solute, G. An example of such a solute is glucose. So the extracellular solute concentration space is $c^{\text{f(b)}}_{i,0} = \{c^{\text{f(b)}}_{\text{Na},0}, c^{\text{f(b)}}_{\text{K},0}, c^{\text{f(b)}}_{\text{Cl},0}, c^{\text{f(b)}}_{\text{H},0}, c^{\text{f(b)}}_{\text{HCO}_3,0}, c^{\text{f(b)}}_{\text{A},0}, c^{\text{f(b)}}_{\text{G},0}\}^T$. Here we allow the extracellular solute concentration to be different at the two ends of the cell.

The $c$ and $c_0$ in Eq. S4 are defined by

$$c = \sum c_i\,, \quad c^{\text{f(b)}}_0 = \sum c^{\text{f(b)}}_{i,0}\,, \tag{S11}$$

which are the osmolarities inside and outside of the cell.

#### 1.6 Chemical reactions and pH

We consider two types of chemical reactions in the model: the bicarbonate-carbonic acid pair and the buffer pair. The chemical equilibrium equation for the bicarbonate-carbonic acid pair is

$$CO_2(\text{aq}) + H_2O(\text{l}) \rightleftharpoons H^+(\text{aq}) + HCO_3^-(\text{aq})\,, \tag{S12}$$

where $[CO_2]_{aq}$ is related to the partial pressure of $CO_2$, $P_{CO_2}$, by the Henry constant $k_H$, $[CO_2]_{aq} = P_{CO_2}/k_H$. The reaction equilibrium constant is

$$k_c = \frac{[HCO_3^-]_{aq}[H^+]_{aq}}{[CO_2]_{aq}} . \tag{S13}$$

We let $pH_0 = -\log_{10}[H^+]_{aq,0}$ as the extracelluar pH and $pK_c = -\log_{10}k_c$ so that Eq. S13 becomes

$$pH_0 - pK_c = \log_{10}\frac{[HCO_3^-]^0_{aq}}{P_{CO_2}/k_H} . \tag{S14}$$

$[CO_2]_{aq} = [CO_2]^0_{aq}$ since $CO_2$ can move freely across the cell membrane [20]. Once the extracellular concentration of bicarbonate and the partial pressure of $CO_2$ are given, the extracellular pH is also uniquely determined. Similarly, in the intracellular domain, we have

$$pH - pK_c = \log_{10}\frac{[HCO_3^-]_{aq}}{[CO_2]_{aq}} , \tag{S15}$$

where $pH = -\log_{10}[H^+]_{aq}$ is the intracelluar pH. The chemical reaction for the intracellular buffer solution is

$$\text{HBuf(aq)} \rightleftharpoons \text{H}^+\text{(aq)} + \text{Buf}^-\text{(aq)} . \tag{S16}$$

The reaction equilibrium constant is similarly $k_B = [\text{Buf}^-]_{aq}[H^+]_{aq}/[\text{HBuf}]_{aq}$. With $pK_B = -\log_{10}k_B$, we have

$$pH - pK_B = \log_{10}\frac{[\text{Buf}^-]_{aq}}{[\text{HBuf}]_{aq}} . \tag{S17}$$

A model with a more involved buffer solution reaction has been considered where the valance of the deprotonated buffer can take on any numbers [21]. However, this will not affect our model prediction.

Since we assume that the buffer solution only exists inside the cell, it is equivalent to saying that the buffer solutes are non-permeable. The total amount of non-permeable solutes is conserved within the cell so that

$$S\int_{x^b}^{x^f} c_A dx = N_A , \quad S\int_{x^b}^{x^f} (c_{\text{Buf}} + c_{\text{HBuf}})dx = N_{\text{Buf}} + N_{\text{HBuf}} , \tag{S18}$$

where $S$ is the cross-sectional area of the cell, $N_A$ is the total amount of intracellular non-permeable large molecules, and $N_{\text{Buf}} + N_{\text{HBuf}}$ is the total amount of buffer solutes inside the cell. Both $N_A$ and $N_{\text{Buf}} + N_{\text{HBuf}}$ are given parameters in the model.

With all the relations established above, we can express the concentrations of hydrogen, bicarbonate, and protonated buffer solute as functions of the primary variables, i.e.,

$$c_{\text{H}}(x) = 10^3 10^{-\text{pH}} , \quad c_{\text{HCO}_3}(x) = \frac{P_{CO_2}}{k_H}10^{\text{pH}-\text{p}K_c} , \quad c_{\text{HBuf}}(x) = c_{\text{Buf}}10^{\text{p}K_B-\text{pH}} , \tag{S19}$$

which should be satisfied at all points in space.

##### 1.7 Ionic dynamics

Diffusion, convection, and electric drift happen for each solute species in the cytosol. The flux for each species is

$$J_i = -D_i \frac{dc_i}{dx} + v_c c_i - D_i \frac{z_i F}{RT} c_i \frac{d\phi}{dx}\,, \quad i \in \{\mathrm{Na}^+, \mathrm{K}^+, \mathrm{Cl}^-, \mathrm{H}^+, \mathrm{HCO}_3^-, \mathrm{A}^-, \mathrm{Buf}^-, \mathrm{HBuf}\}\,, \tag{S20}$$

where $c_i$, $z_i$, and $D_i$ are, respectively, each solute species' concentration, valance, and diffusion constant (note that $z_{\mathrm{HBuf}} = 0$). $\phi$ is the intracellular electric potential. $F$, $R$, and $T$ are Faraday's constant, ideal gas constant, and absolute temperature, respectively.

The governing equations and boundary conditions for $c_{\mathrm{Na}}$, $c_{\mathrm{K}}$, $c_{\mathrm{Cl}}$, and $c_{\mathrm{A}}$, which are non-reactive species, are

$$-\frac{dJ_{\mathrm{Na}}}{dx} = 0\,, \quad J_{\mathrm{Na}}\big|_{x=x^{\mathrm{f}}} = -J^{\mathrm{f}}_{\mathrm{Na}}\,, \quad J_{\mathrm{Na}}\big|_{x=x^{\mathrm{b}}} = J^{\mathrm{b}}_{\mathrm{Na}}\,, \tag{S21}$$

$$-\frac{dJ_{\mathrm{K}}}{dx} = 0\,, \quad J_{\mathrm{K}}\big|_{x=x^{\mathrm{f}}} = -J^{\mathrm{f}}_{\mathrm{K}}\,, \quad J_{\mathrm{K}}\big|_{x=x^{\mathrm{b}}} = J^{\mathrm{b}}_{\mathrm{K}}\,, \tag{S22}$$

$$-\frac{dJ_{\mathrm{Cl}}}{dx} = 0\,, \quad J_{\mathrm{Cl}}\big|_{x=x^{\mathrm{f}}} = -J^{\mathrm{f}}_{\mathrm{Cl}}\,, \quad J_{\mathrm{Cl}}\big|_{x=x^{\mathrm{b}}} = J^{\mathrm{b}}_{\mathrm{Cl}}\,, \tag{S23}$$

$$-\frac{dJ_{\mathrm{A}}}{dx} = 0\,, \quad J_{\mathrm{A}}\big|_{x=x^{\mathrm{f}}} = J_{\mathrm{A}}\big|_{x=x^{\mathrm{b}}} = 0\,, \tag{S24}$$

where the boundary fluxes are positive inwards. The governing equation and boundary conditions for pH, i.e., $c_{\mathrm{H}}$, are

$$\begin{aligned} &-\frac{d}{dx}\left(J_{\mathrm{HCO_3}} + J_{\mathrm{Buf}} - J_{\mathrm{H}}\right) = 0\,, \\ &\left(J_{\mathrm{HCO_3}} + J_{\mathrm{Buf}} - J_{\mathrm{H}}\right)\big|_{x=x^{\mathrm{f}}} = -\left(J^{\mathrm{f}}_{\mathrm{HCO_3}} + J^{\mathrm{f}}_{\mathrm{Buf}} - J^{\mathrm{f}}_{\mathrm{H}}\right)\,, \\ &\left(J_{\mathrm{HCO_3}} + J_{\mathrm{Buf}} - J_{\mathrm{H}}\right)\big|_{x=x^{\mathrm{b}}} = \left(J^{\mathrm{b}}_{\mathrm{HCO_3}} + J^{\mathrm{b}}_{\mathrm{Buf}} - J^{\mathrm{b}}_{\mathrm{H}}\right)\,. \end{aligned} \tag{S25}$$

where $J^{\mathrm{b/f}}_{\mathrm{Buf}} = 0$ due to the assumed non-permeability of buffer solutions. This governing equation for pH results from Eqs. S12 and S16. The governing equation and boundary conditions for $c_{\mathrm{Buf}}$ are

$$-\frac{d}{dx}\left(J_{\mathrm{Buf}} + J_{\mathrm{HBuf}}\right) = 0\,, \quad \left(J_{\mathrm{Buf}} + J_{\mathrm{HBuf}}\right)\big|_{x=x^{\mathrm{f}}} = \left(J_{\mathrm{Buf}} + J_{\mathrm{HBuf}}\right)\big|_{x=x^{\mathrm{b}}} = 0\,. \tag{S26}$$

The intracellular electric potential is solved by the electro-neutrality condition, i.e.,

$$\sum z_i c_i = 0\,. \tag{S27}$$

The same condition is also imposed outside the cell when we set up the parameters for extracellular solute concentrations. This electro-neutrality condition enforces that the net charge is 0 everywhere, leading to $\nabla^2\phi = 0$. We thus know that the intracellular voltage is a linear function in space, i.e., $\nabla\phi = \mathrm{const}$. Therefore, Eq. S20 can be written as

$$J_i = -D_i \frac{dc_i}{dx} + \left(v_c - D_i \frac{z_i F}{RT}\frac{d\phi}{dx}\right) c_i\,. \tag{S28}$$

The term in the parentheses is a constant, meaning that the combined cytosol flow and electric drift generate an effective convection velocity for each solute species.

Note that the ionic dynamics described in this section is generic, meaning that these are intracellular fluxes and are independent of the choice of membrane ion channels, transporters, and pumps, which we discuss next.

#### 1.8 Ion channels, transporters, and pumps

To specify the boundary ion fluxes for each species, $J_i^{\mathrm{b/f}}$, the ion channels, transporters, and pumps need to be modeled. In this work, we choose the following membrane carriers: passive $Na^+$ channel, passive $K^+$ channel, passive $Cl^-$ channel (including SWELL1), $Na^+/K^+$ pump (NKE), $Na^+/H^+$ exchanger (NHE), and $Cl^-/HCO_3^-$ exchanger (AE2).

The passive channels are typically tension-gated [22]. We let $G_m^{\mathrm{f(b)}} \in (0,1)$ be a mechanosensitive gating function that follows a Boltzmann distribution, i.e., $G_m^{\mathrm{f(b)}} = \left[1 + e^{-\beta_\tau(\tau_m^{\mathrm{f(b)}} - \tau_{m,c})}\right]^{-1}$, where $\beta_\tau$ and $\tau_{m,c}$ are two constants and $\tau_m^{\mathrm{f(b)}}$ is the cortical/membrane tension on either end of the cell. The cortical tension can be calculated from the force balance at the membrane, i.e.,

$$\tau_m^{\mathrm{f(b)}} = \frac{b}{2}\left(\sigma^{\mathrm{f(b)}} + p_c^{\mathrm{f(b)}} + p_*^{\mathrm{f(b)}}\right) . \tag{S29}$$

The passive ion fluxes, $J_{i,p}^{\mathrm{f(b)}}$, are proportional to the electrochemical potential difference of ions across the membrane [23],

$$J_{i,p}^{\mathrm{f(b)}} = \alpha_{i,p}^{\mathrm{f(b)}} G_m^{\mathrm{f(b)}} \left[RT \ln \Gamma_i^{\mathrm{f(b)}} - z_i F(\phi^{\mathrm{f(b)}} - \phi_0^{\mathrm{f(b)}})\right] , \quad i \in \{\mathrm{Na}^+, \mathrm{K}^+, \mathrm{Cl}^-\} \tag{S30}$$

where $\Gamma_i^{\mathrm{f(b)}} = c_{i,0}^{\mathrm{f(b)}}/c_i^{\mathrm{f(b)}}$ is the ratio of extra- to intra-cellular ion concentrations at the two ends of the cell; $\alpha_{i,p}^{\mathrm{f(b)}}$ is the permeability coefficient of each species, which depends on the channel property and the density of the channels in the membrane.

The $Na^+/K^+$ pump (NKE) is a ubiquitous and vital active ion pump that maintains the membrane potential of cells. It exports three $Na^+$ ions and intakes two $K^+$ ions per ATP molecule. Because the overall flux is positive outwards, the pump's activity depends on the membrane potential [24]. The NKE flux also depends on the concentrations of $Na^+$ and $K^+$ [25, 26] and saturates at high concentration limits [26]. Based on these facts, we model the flux of $Na^+$ and $K^+$ through the $Na^+/K^+$ pump as

$$J_{\mathrm{NKE}}^{\mathrm{f(b)}} = J_{\mathrm{NKE,Na}}^{\mathrm{f(b)}} = -\frac{3}{2} J_{\mathrm{NKE,K}}^{\mathrm{f(b)}} = -\frac{\alpha_{\mathrm{NKE}}^{\mathrm{f(b)}} G_{V,\mathrm{NKE}}^{\mathrm{f(b)}}}{\left(1 + \beta_{\mathrm{NKE,Na}} \Gamma_{\mathrm{Na}}^{\mathrm{f(b)}}\right)^3 \left(1 + \beta_{\mathrm{NKE,K}}/\Gamma_{\mathrm{K}}^{\mathrm{f(b)}}\right)^2} , \tag{S31}$$

where $\alpha_{\mathrm{NKE}}^{\mathrm{f(b)}}$ is the permeability coefficient of the pump depending on the density of the pump as well as the concentration of ATP. $\beta_{\mathrm{NKE,Na}}$ and $\beta_{\mathrm{NKE,K}}$ are constants that scale $\Gamma_{\mathrm{Na}}^{\mathrm{f(b)}}$ and $\Gamma_{\mathrm{K}}^{\mathrm{f(b)}}$, respectively. These two constants are intrinsic properties of the NKE pump and, thus, do not differ at the two ends of the cells. The exponents 3 and 2 are Hill's coefficients of $Na^+$ and $K^+$, respectively. Equation S31 ensures that the flux is zero when either $1/\Gamma_{\mathrm{Na}}^{\mathrm{f(b)}}$ or $\Gamma_{\mathrm{K}}^{\mathrm{f(b)}}$ approaches zero; the flux saturates if $1/\Gamma_{\mathrm{Na}}^{\mathrm{f(b)}}$ and $\Gamma_{\mathrm{K}}^{\mathrm{f(b)}}$ approaches infinity. $G_{V,\mathrm{NKE}}^{\mathrm{f(b)}}$ captures the voltage-dependence of the pump activity [24], $G_{V,\mathrm{NKE}}^{\mathrm{f(b)}} = 2\left[1+e^{-\beta_\phi\left(\phi^{\mathrm{f(b)}}-\phi_c\right)}\right]^{-1} - 1$, where $\beta_\phi$ and $\phi_c$ are constants that reflects the intrinsic voltage-gating property of NKE.

The $Na^+/H^+$ exchanger (NHE), which has ten identified isoforms, is expressed in almost all tissues [20]. It imports one $Na^+$ and extrudes one $H^+$ under physiological conditions. This exchanger has significant effects on water flux, cell volume regulation [27–32], cell migration [2, 33–35], and is a major therapeutic target [20, 36, 37]. NHE is quiescent at intracellular pH $> 7.2$ [38]. The flux of NHE can thus be expressed as

$$J_{\mathrm{NHE}}^{\mathrm{f(b)}} = J_{\mathrm{NHE,Na}}^{\mathrm{f(b)}} = -J_{\mathrm{NHE,H}}^{\mathrm{f(b)}} = \alpha_{\mathrm{NHE}}^{\mathrm{f(b)}} G_{\mathrm{NHE}}^{\mathrm{f(b)}} RT\left(\ln\Gamma_{\mathrm{Na}}^{\mathrm{f(b)}} - \ln\Gamma_{\mathrm{H}}^{\mathrm{f(b)}}\right), \tag{S32}$$

where $\alpha_{\mathrm{NHE}}^{\mathrm{f(b)}}$ is the permeability coefficient which does not significantly depend on cortical tension [39]. However, $\alpha_{\mathrm{NHE}}^{\mathrm{f(b)}}$ depends on the density of membrane NHE, which is affected by multiple factors, including the presence of F-actin on the cell membrane [4,20]. $G_{\mathrm{NHE}}^{\mathrm{f(b)}} = \left[1+e^{\beta_{\mathrm{NHE}}\left(\mathrm{pH}^{\mathrm{f(b)}}-\mathrm{pH}_{\mathrm{NHE},c}\right)}\right]^{-1}$ is a pH-gated function indicating the dependence of the NHE activity on pH. $\beta_{\mathrm{NHE}}$ and $\mathrm{pH}_{\mathrm{NHE},c}$ are two constants reflecting the NHE properties.

The $Cl^-/HCO_3^-$ exchanger (AE2), which imports one $Cl^-$ and extrudes one $HCO_3^-$, is also common in cells. This exchanger is almost quiescent at intracellular pH $< 6.8 - 7.3$. Similarly, we assume that the flux takes the form

$$J_{\mathrm{AE2}}^{\mathrm{f(b)}} = J_{\mathrm{AE2,Cl}}^{\mathrm{f(b)}} = -J_{\mathrm{AE2,HCO_3}}^{\mathrm{f(b)}} = \alpha_{\mathrm{AE2}}^{\mathrm{f(b)}} G_{\mathrm{AE2}}^{\mathrm{f(b)}} RT\left(\ln\Gamma_{\mathrm{Cl}}^{\mathrm{f(b)}} - \ln\Gamma_{\mathrm{HCO_3}}^{\mathrm{f(b)}}\right), \tag{S33}$$

where $\alpha_{\mathrm{AE2}}^{\mathrm{f(b)}}$ is the permeability coefficient of AE2 and is assumed to be independent of the cortical tension. $G_{\mathrm{AE2}}^{\mathrm{f(b)}} = \left[1+e^{-\beta_{\mathrm{AE2}}\left(\mathrm{pH}^{\mathrm{f(b)}}-\mathrm{pH}_{\mathrm{AE2},c}\right)}\right]^{-1}$ is a pH-gated function indicating the dependence of the AE2 activity on pH. $\beta_{\mathrm{AE2}}$ and $\mathrm{pH}_{\mathrm{AE2},c}$ are two constants reflecting the AE2 properties.

With all these ion carriers considered, the boundary fluxes for each ionic species are:

$$J_{\mathrm{Na}}^{\mathrm{f(b)}} = J_{\mathrm{Na},p}^{\mathrm{f(b)}} + J_{\mathrm{NKE,Na}}^{\mathrm{f(b)}} + J_{\mathrm{NHE,Na}}^{\mathrm{f(b)}}, \tag{S34}$$

$$J_{\mathrm{K}}^{\mathrm{f(b)}} = J_{\mathrm{K},p}^{\mathrm{f(b)}} + J_{\mathrm{NKE,K}}^{\mathrm{f(b)}}, \tag{S35}$$

$$J_{\mathrm{Cl}}^{\mathrm{f(b)}} = J_{\mathrm{Cl},p}^{\mathrm{f(b)}} + J_{\mathrm{AE2,Cl}}^{\mathrm{f(b)}}, \tag{S36}$$

$$J_{\mathrm{H}}^{\mathrm{f(b)}} = J_{\mathrm{NHE,H}}^{\mathrm{f(b)}}, \tag{S37}$$

$$J_{\mathrm{HCO_3}}^{\mathrm{f(b)}} = J_{\mathrm{AE2,HCO_3}}^{\mathrm{f(b)}}. \tag{S38}$$

#### 1.9 Force balance of the cell

At the back of the cell, F-actin adheres to the substrate through transmembrane proteins (integrins), which provide an adhesive force, $F_{\mathrm{ad}}^{\mathrm{b}}$, that resists cell migration. We let the adhesive force be proportional to the cell velocity, i.e., $F_{\mathrm{ad}}^{\mathrm{b}} = k_{\mathrm{ad}} v_{\mathrm{cell}}$, where $k_{\mathrm{ad}}$ is the coefficient of adhesive force. This adhesive force is equivalent to the effective frictional force between the cell and the substrate.

To establish a force balance relation of the cell, we can draw a free-body diagram of the cell and collect all the forces applied to the cell either from the extracellular matrix or through other external means. Any internal forces within the cell should be excluded when analyzing a free-body diagram. This condition also means the choices of the constitutive relations for the actin network, such as actin swelling and myosin contraction, do not affect the force balance of the cell.

Putting together, the force balance of the entire cell is

$$-(p_*^{\mathrm{f}} - p_*^{\mathrm{b}}) - \eta_{\mathrm{st}} \int_{x^{\mathrm{b}}}^{x^{\mathrm{f}}} \theta_n v_n dx - k_{\mathrm{ad}} v_{\mathrm{cell}} = 0\,, \tag{S39}$$

where $p_*$'s are defined in Eq. S5.

The primary variables (Eq. S1) are numerically solved by solving all the coupled nonlinear equations and boundary conditions. The one-dimensional space is discretized into elements where the Finite Difference Method is applied. The discretized equations are solved with the Newton–Raphson iteration scheme. Other variables are expressed in terms of the unknowns or are considered model parameters.

##### 1.10 Coupling between mechanics and chemistry

The model presented above is tightly coupled. None of the equations can be solved without solving all the other equations together. We now extend the basic model to include additional couplings between the mechanical and biochemical parts of the model. Specifically, we let pH and NHE be coupled with the actin network. This effort is achieved through two parallel schemes.

The rate of actin depolymerization, $\gamma$, depends on multiple mechanical and biochemical factors [40]. One of the factors we are considering here is pH [41]. Experimental data suggests that higher pH leads to a higher rate of actin depolymerization [41]. We thus let

$$\gamma = \gamma_0 + \frac{\gamma_{\mathrm{pH}}}{1 + e^{-\beta_\gamma(\mathrm{pH} - \mathrm{pH}_{\gamma,c})}}\,, \tag{S40}$$

where $\gamma_0$ is a baseline rate of actin depolymerization, and $\gamma_{\mathrm{pH}}$ is the coefficient of pH-dependent rate of actin depolymerization. $\beta_\gamma$ and $\mathrm{pH}_{\gamma,c}$ are two constants characterizing how $\gamma$ depends on pH. Note that pH is a function of space $x$, so $\gamma$ is also a spatial function.

Our recent work showed that the expression of NHE is coupled to F-actin density [4], a mechanism controlled by a protein known as ezrin [20]. In this work, we will study, from a theoretical point of view, how this coupling affects cell homeostasis and migration. Since the membrane expression of NHE decreases with decreasing F-actin presence, we let the permeability coefficient of NHE be a function of F-actin concentration

$$\alpha_{\mathrm{NHE}}^{\mathrm{f(b)}} = \frac{\alpha_{\mathrm{NHE,F}}^{\mathrm{f(b)}}}{1 + e^{-\beta_F\left(\theta_n^{\mathrm{f(b)}}/\theta_{n,*} - \beta_{F,c}\right)}}\,, \tag{S41}$$

where $\alpha_{\mathrm{NHE,F}}^{\mathrm{f(b)}}$ is the maximum permeability coefficient, which depends on cell types and other transcription factors. $\beta_F$ and $\beta_{F,c}$ are constants. When the actin-dependence of NHE permeability is muted, we will let $\alpha_{\mathrm{NHE}}^{\mathrm{f(b)}} = \alpha_{\mathrm{NHE,F}}^{\mathrm{f(b)}} = $ conts., which can be different at the two ends of the cell.

#### 2 Model Parameters

All the parameters used in the model are listed in Tabs. S1 and S2, which represent a typical mammalian tissue cell under physiological conditions. When we show results from polarization ion carriers, we specify the polarization ratio as the ratio of ion permeability at the front to the rate at the back. The average permeability will remain the same as those listed in Tab. S2 while the front and back values are adjusted accordingly, i.e., $(\alpha^{\mathrm{f}} + \alpha^{\mathrm{b}})/2 = \alpha$ is fixed.

Table S1: Model parameters on the mechanical part.

| Parameter | Description | Value | Source |
|---|---|---|---|
| $R$ (J/mol/K) | Ideal gas constant | 8.31 | Physical constant |
| $T$ (K) | Absolute temperature | 310 | Physiological condition |
| $L$ (μm) | Cell length | 50 | Based on Ref. [3] |
| $b$ (μm) | Cell width | 3 | Based on Ref. [3] |
| $w$ (μm) | Cell depth | 10 | Based on Ref. [3] |
| $\eta$ (Pa·s/μm$^2$/mM ) | Drag coefficient between two phases | $10^{-2}$ | Based on Ref. [42] |
| $\eta_{\mathrm{st}}$ (Pa·s/μm$^2$/mM) | Coefficient of drag from focal adhesion | $10^{2}$ | Based on Refs. [3, 11] |
| $k_{\sigma_n}$ (Pa/mM) | Coefficient of the passive F-actin stress | $10^{2}$ | Based on Refs. [3, 11] |
| $k_{\mathrm{ad}}$ (Pa·s/μm) | Coefficient in $F_{\mathrm{ad}}^{\mathrm{b}} = k_{\mathrm{ad}} v_{\mathrm{cell}}$ | 300 | Based on Refs. [3, 11] |
| $d_g$ (Pa·s/μm) | Coefficient of hydraulic pressure | 20 | Ref. [3] |
| $D_{\theta_c}$ (μm$^2$/s) | Diffusion coefficient of G-actin | 10 | Refs. [43–46] |
| $\theta_*$ (mM) | Average concentration of total actin | 0.3 | Based on Ref. [47, 48] |
| $J_a$ (nm mM/s) | Coefficient in $J_{\mathrm{actin}}^{\mathrm{f}} = J_a \theta_c^{\mathrm{f}} / (\theta_{c,c} + \theta_c^{\mathrm{f}})$ | 6 | Ref. [11] |
| $\theta_{c,c}$ (μM) | Critical value of actin polymerization | 0.2 | Ref. [48] |
| $\gamma_0$ (1/s) | Baseline rate of actin depolymerization | $10^{-3}$ | Based on Ref. [11] |
| $\gamma_{\mathrm{pH}}$ (1/s) | Coefficient of pH-dependent rate of actin depolymerization | $2 \times 10^{-3}$ | Estimated |
| $\beta_\gamma$ | Constant in $\gamma$ | 15 | Estimated |
| $\mathrm{pH}_{\gamma,c}$ | Constant in $\gamma$ | 7.38 | Estimated |
| $p_\infty$ (Pa) | External pressure at the front and back | 0 | Free parameter |

Table S2: Model parameters on the electrochemical part.

| Parameter | Description | Value | Source |
|---|---|---|---|
| $N_A$ (pmol) | Total intracellular $A^-$ | $7 \times 10^{-2}$ | Based on Ref. [3] |
| $N_{\mathrm{Buf}} + N_{\mathrm{HBuf}}$ (pmol) | Total intracellular Buffer Solution | $7 \times 10^{-2}$ | Based on Ref. [3] |
| $\alpha_{\mathrm{w}}$ (nm/Pa/s) | Permeability coefficient of water | 0.1 | Ref. [49] |
| $\alpha_{\mathrm{Na},p}$ ($\mathrm{mol}^2/\mathrm{J}/\mu\mathrm{m}^2/\mathrm{s}$) | Permeability of Na | 0.1 | Based on Refs. [3, 21] |
| $\alpha_{\mathrm{K},p}$ ($\mathrm{mol}^2/\mathrm{J}/\mu\mathrm{m}^2/\mathrm{s}$) | Permeability of K | $1.5 \times 10^2$ | Based on Refs. [3, 21] |
| $\alpha_{\mathrm{Cl},p}$ ($\mathrm{mol}^2/\mathrm{J}/\mu\mathrm{m}^2/\mathrm{s}$) | Permeability of Cl | $10^2$ | Based on Refs. [3, 21] |
| $\alpha_{\mathrm{NKE}}$ ($\mathrm{mol}/\mu\mathrm{m}^2/\mathrm{s}$) | Permeability of NKE | $3.9 \times 10^3$ | Based on Refs. [3, 21] |
| $\alpha_{\mathrm{NHE,F}}$ ($\mathrm{mol}^2/\mathrm{J}/\mu\mathrm{m}^2/\mathrm{s}$) | Permeability of NHE | $10^3$ | Based on Refs. [3, 21] |
| $\alpha_{\mathrm{AE2}}$ ($\mathrm{mol}^2/\mathrm{J}/\mu\mathrm{m}^2/\mathrm{s}$) | Permeability of AE2 | $5 \times 10^2$ | Based on Refs. [3, 21] |
| $\beta_{\mathrm{NKE,Na}}$ | Constant in $J_{\mathrm{NKE}}$ | 0.1 | Based on Refs. [3, 21] |
| $\beta_{\mathrm{NKE,K}}$ | Constant in $J_{\mathrm{NKE}}$ | 0.01 | Based on Refs. [3, 21] |
| $\beta_\tau$ (m/N) | Constant in $G_m$ | $2 \times 10^3$ | Based on Refs. [3, 21] |
| $\tau_{m,c}$ (N/m) | Constant in $G_m$ | $5 \times 10^{-4}$ | Based on Refs. [3, 21] |
| $\beta_\phi$ (1/mV) | Constant in $G_{V,\mathrm{NKE}}$ | 0.03 | Ref. [24] |
| $\phi_c$ (mV) | Constant in $G_{V,\mathrm{NKE}}$ | $-150$ | Ref. [24] |
| $\beta_{\mathrm{NHE}}$ | Constant in $G_{\mathrm{NHE}}$ | 15 | Ref. [38] |
| $\mathrm{pH}_{\mathrm{NHE},c}$ | Constant in $G_{\mathrm{NHE}}$ | 7.2 | Ref. [38] |
| $\beta_{\mathrm{AE2}}$ | Constant in $G_{\mathrm{AE2}}$ | 10 | Ref. [38] |
| $\mathrm{pH}_{\mathrm{AE2},c}$ | Constant in $G_{\mathrm{AE2}}$ | 7.1 | Ref. [38] |
| $\beta_F$ | Constant in $\alpha_{\mathrm{NHE}}$ | 5 | Estimated |
| $\beta_{F,c}$ | Constant in $\alpha_{\mathrm{NHE}}$ | 0.1 | Estimated |
| $k_H$ (atm/M) | Henry's constant | 29 | Ref. [50] |
| $P_{\mathrm{CO_2}}$ (atm) | Partial pressure of $CO_2$ | 5% | Physiological condition |
| $\mathrm{p}K_c$ | $\mathrm{p}K$ for bicarbonate-carbonic acid pair | 6.1 | Ref. [50] |
| $\mathrm{p}K_B$ | $\mathrm{p}K$ for intracellular buffer | 7.5 | Based on Ref. [50] |
| $c_{\mathrm{Na},0}$ (mM) | $Na^+$ concentration in the medium | 145 | Physiological condition |
| $c_{\mathrm{K},0}$ (mM) | $K^+$ concentration in the medium | 9 | Physiological condition |
| $c_{\mathrm{Cl},0}$ (mM) | $Cl^-$ concentration in the medium | 105 | Physiological condition |
| $c_{\mathrm{HCO_3},0}$ (mM) | $HCO_3^-$ concentration in the medium | 35 | Physiological condition |
| $c_{\mathrm{G},0}$ (mM) | Glucose concentration in the medium | 25 | Physiological condition |

#### Acknowledgment

Yizeng Li is supported by NSF 2303648. Sean X Sun is supported by NIH R01GM134542. The opinions, findings, and conclusions, or recommendations expressed are those of the authors and do not necessarily reflect the views of any of the funding agencies.

#### Supplementary Figures

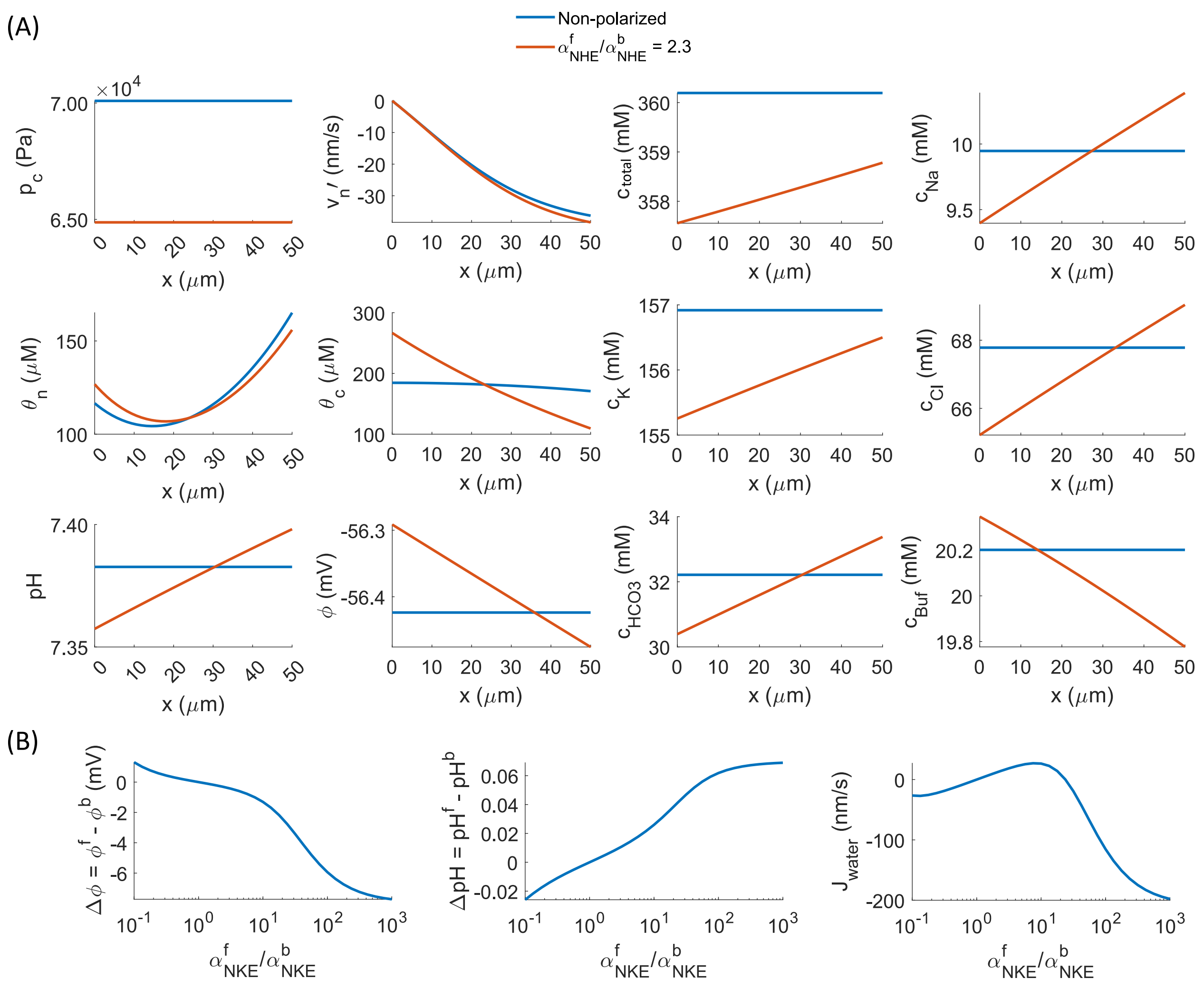


Figure S1: Model prediction from a cell without coupling between mechanics and biochemistry. (A) Spatial distribution of field variables from a non-polarized cell and a cell with front polarization of NHE. $p_c$: intracellular cytosol pressure. $v_n'$: F-actin velocity in the frame of the cell. $\theta_n$: F-actin concentration. $\theta_c$: G-actin concentration. $\phi$: intracellular electric potential. $c_{\text{total}}$: total intracellular solute concentration. (B) Intracellular pH difference ($\Delta$pH), electric potential difference ($\Delta\phi$), and water flux as functions of the polarization ratio of NKE.

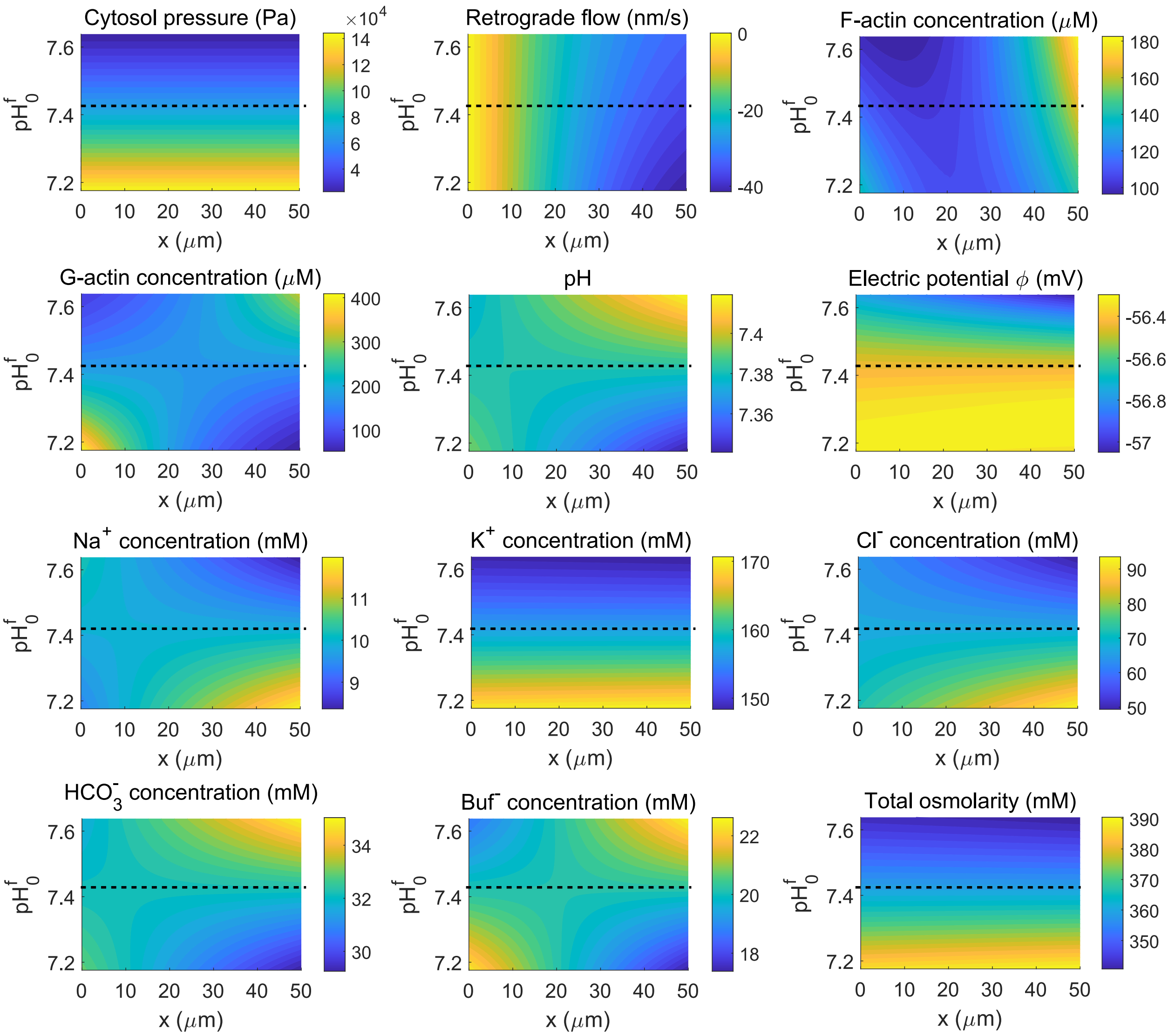


Figure S2: Spatial distribution of variables as the extracellular pH varies at the cell front. Extracellular pH change is achieved by changing the concentration of bicarbonate. Before polarization, $pH_0^f = pH_0^b = 7.42$. The dashed lines represent at non-polarized position at $pH_0^f = 7.42$. No coupling between mechanics and chemistry.

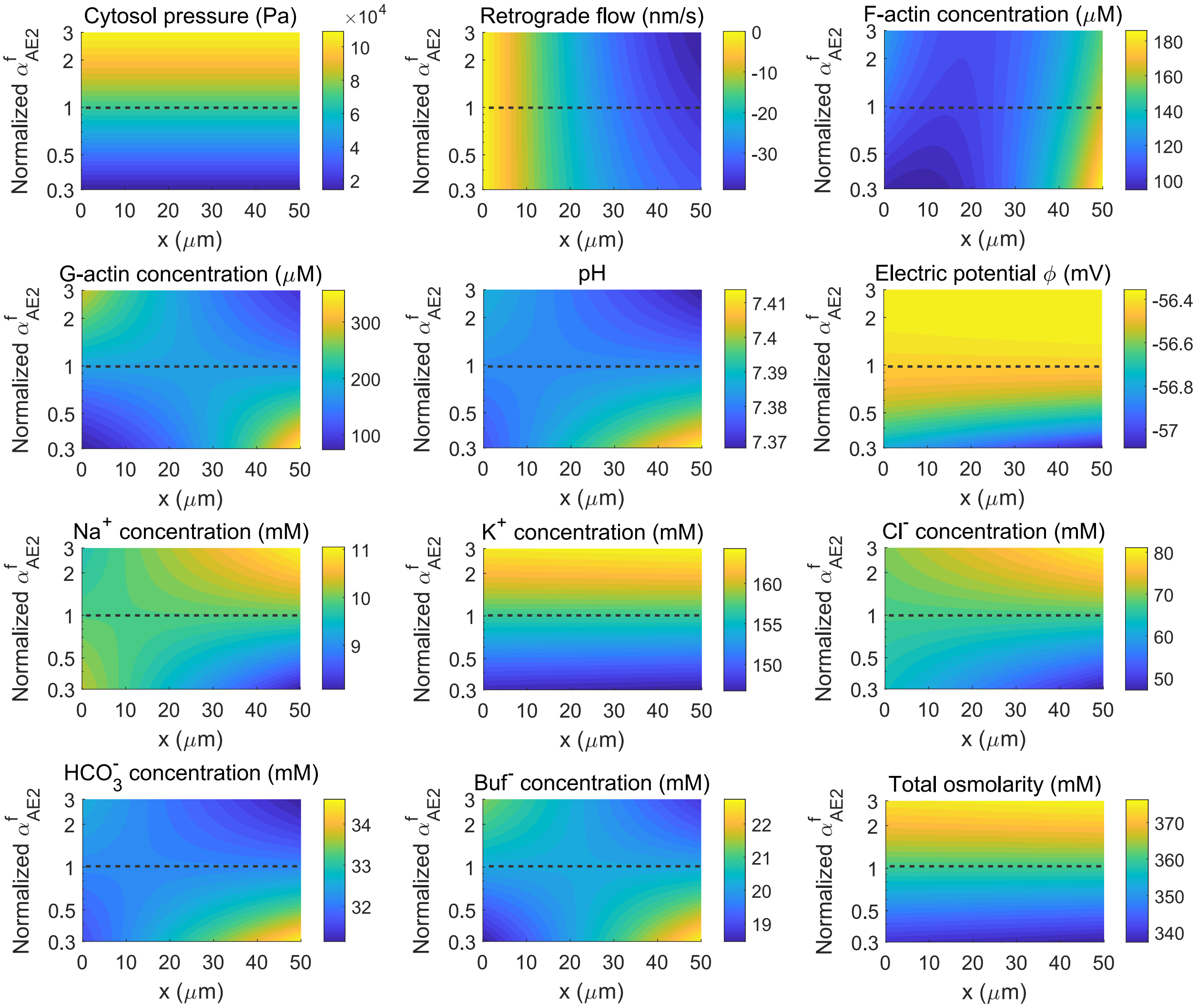


Figure S3: Spatial distribution of variables as the permeability of AE2 varies at the cell front. The coefficient of AE2 at the back is kept constant as provided in the parameter table. The value of the AE2 at the front is normalized with respect to the value in the parameter table. A normalized value less than 1 means the permeability of AE2 at the front is reduced. The dashed lines represent at non-polarized position where $\alpha^{\rm f}_{\rm AE2} = \alpha^{\rm b}_{\rm AE2}$. No coupling between mechanics and chemistry.

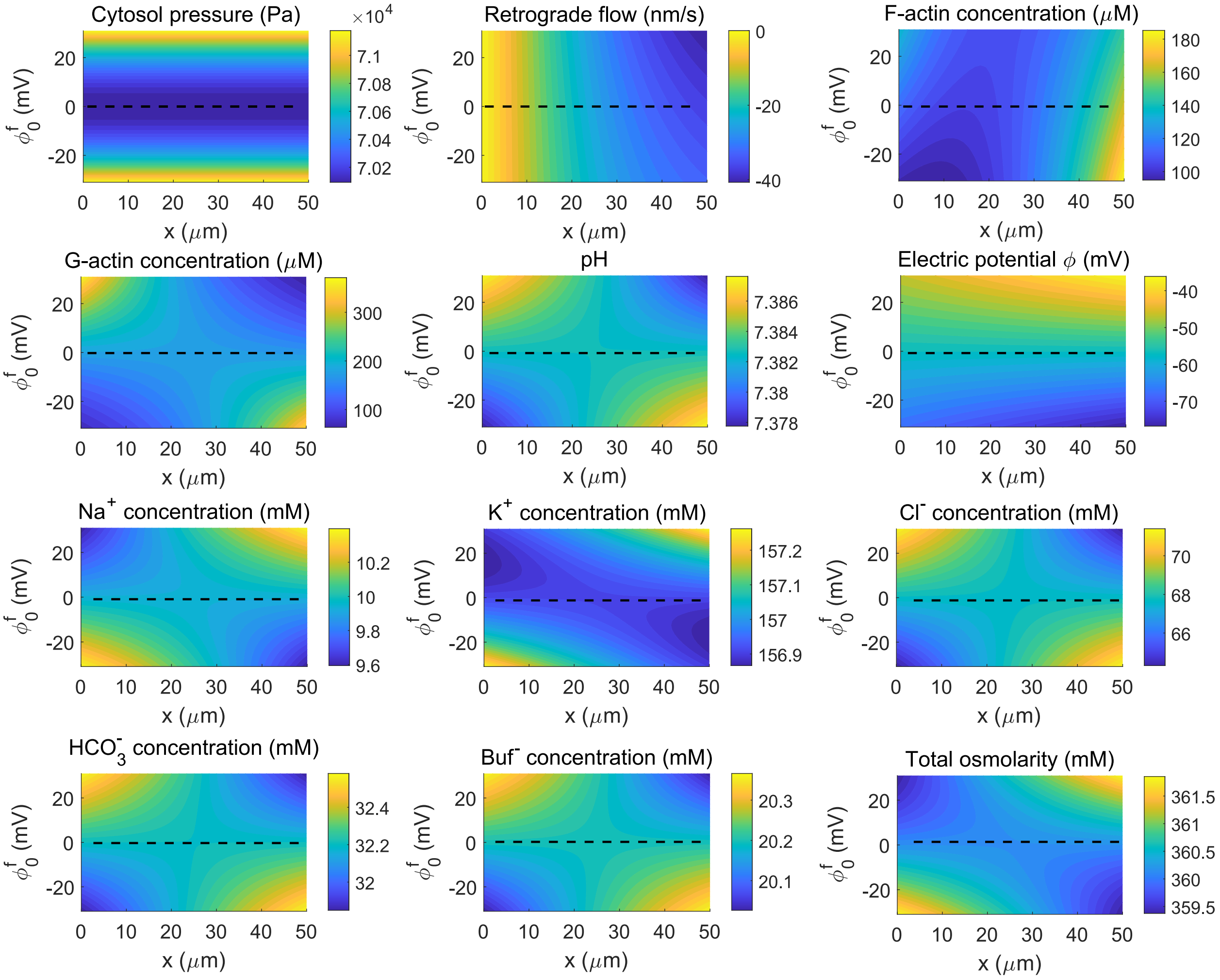


Figure S4: Spatial distribution of variables as the extracellular electric potential varies at the cell front. The dashed lines represent non-polarized position at $\phi_0^{\mathrm{f}} = \phi_0^{\mathrm{b}} = 0$. No coupling between mechanics and chemistry.

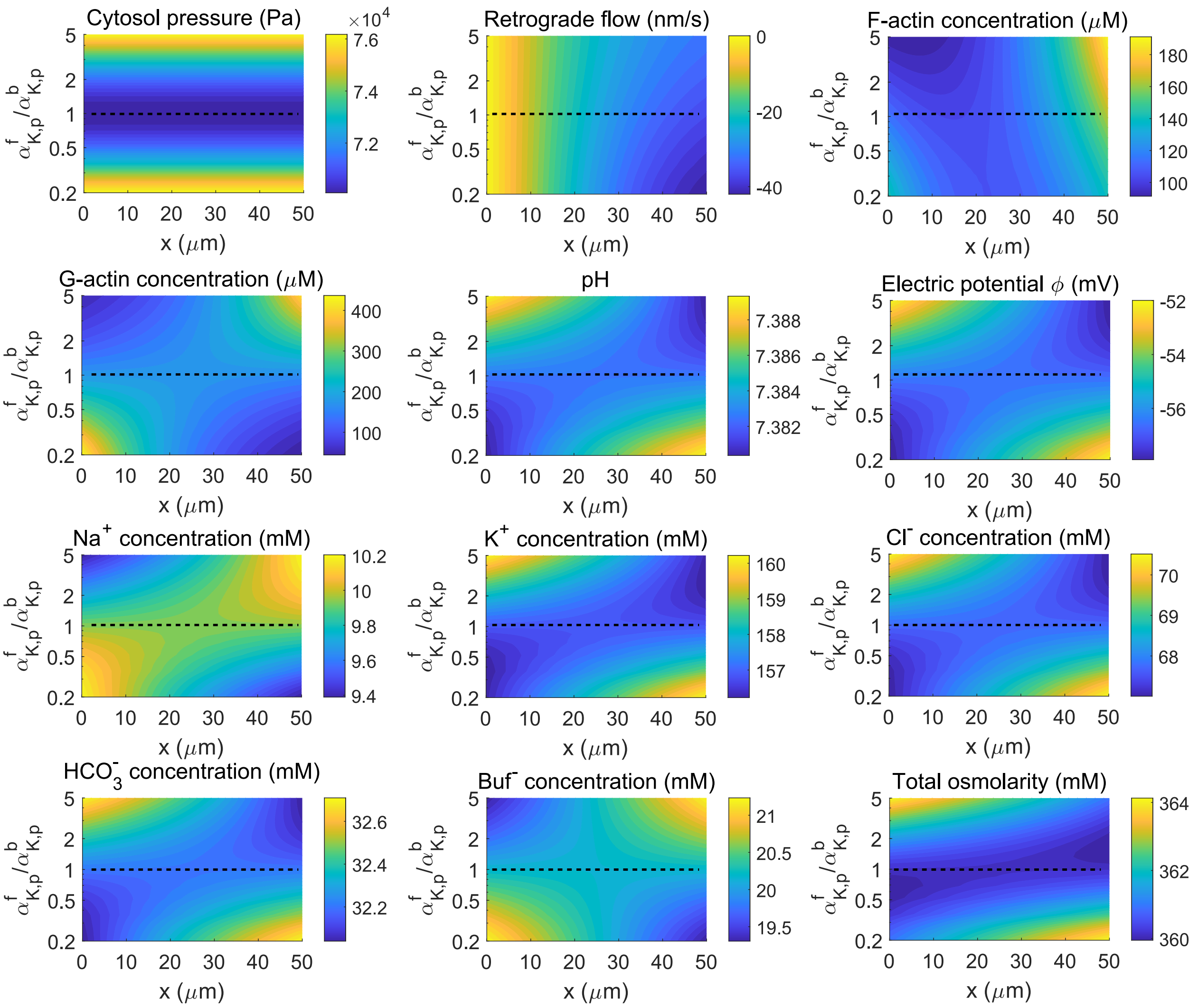


Figure S5: Spatial distribution of variables as the permeability of the passive potassium channel varies at the cell front. The coefficient of the passive potassium channel at the back is kept constant as provided in the parameter table. The value of the passive potassium channel at the front is normalized with respect to the value in the parameter table. A normalized value less than 1 means the permeability of the passive potassium channel at the front is reduced. The dashed lines represent at non-polarized position where $\alpha^{\mathrm{f}}_{\mathrm{K,p}} = \alpha^{\mathrm{b}}_{\mathrm{K,p}}$. No coupling between mechanics and chemistry.
